# A new class of proteinaceous biomaterial with unprecedented properties in unicellular eukaryotes

**DOI:** 10.64898/2025.12.22.695910

**Authors:** Maximilian H. Ganser, Markus Wiederstein, Christof Regl, Laura A. Katz, Sabine Agatha

## Abstract

Biomaterials fundamentally reshape material use, providing superior properties and sustainable alternatives relevant to medicine, textiles, and high-tech applications. Research mainly focuses on animal-derived proteinaceous biomaterials, which remain challenging to reproduce while retaining their remarkable properties. We discovered that the shells of tintinnid ciliates, a group of planktonic unicellular eukaryotes, are composed of self-assembling structural proteins. The shells form in water, are structurally diverse, and show exceptional resistance against high temperatures and the strongest chemicals. Combining single-cell transcriptomics and mass-spectrometry of the shells, we identified the amino acid sequences of the shell-forming proteins. They are exceptionally rich in aromatic residues and possess a unique architecture with flexible, unfolded segments connecting highly stable beta-sheets. This easily accessible system promises new aspects to advance current biomaterial design.

## Main Text

Structural proteins are fundamental building blocks for diverse biological systems, occurring in all lineages across the tree of life and in viruses. Their primary functions include maintaining cellular shape and integrity, enabling movement, and forming protective barriers. Beyond these ubiquitous roles, structural proteins forming complex biomaterials with exceptional mechanical and functional properties have evolved in diverse organisms (*1*). A well-known example is silk, a natural biomaterial secreted by various arthropods, including insects and especially spiders, which is renowned for its high tensile strength, elasticity, and biocompatibility (*2, 3*).

Human use of biomaterials has substantially increased with recent technological progress enabling researchers to mimic various natural properties based on decades of interdisciplinary research (*4*). Applications of the resulting synthetic biomaterials are versatile across industries and include textiles (*5*), cosmetics, coatings, and wet adhesives (*6*). In the medical field, biomaterials are used for tissue engineering, wound dressings, implants, surgical sutures, and drug delivery systems (*7*). More broadly, renewable, biodegradable, and sustainable biomaterials offer environmental benefits and economic potential, as demand for green alternatives to petroleum-based materials like conventional plastics continues to grow (*8*). While many biomaterials have been discovered in Metazoa (*1*), unicellular eukaryotes that comprise the vast majority of eukaryotic diversity (*9*) have been largely overlooked.

Tintinnids are microscopic shell-forming ciliates usually 50–400 µm long that have inhabited the marine plankton since about 260 mya (*10*). Each of the about 1,000 extant species distributed across the world’s oceans and freshwater bodies forms a distinct vase-or tube-shaped shell (lorica) (*11, 12*). Tintinnid shells are intricate structures, rich in texture and ornamentation (*11, 13*), resembling delicate works of art, yet crafted by a single cell (Fig. 1). These shells may be agglutinated with foreign particles (e.g., *Tintinnopsis* species), entirely transparent (e.g., *Schmidingerella* species), or a combination of both (e.g., *Codonellopsis* and *Stenosemella* species). The shell walls can be soft (e.g., *Tintinnidium* and *Antetintinnidium* species) or rigid (remaining tintinnids) and may feature minute pores (e.g., *Schmidingerella*) or large fenestrations (e.g., *Dictyocysta*). At the ultrastructural level, the shell walls consist of one or multiple layers, which exhibit various organizational patterns including chambered, tubular, solid, or crystalline textures (*13*). Previous experiments demonstrated the resilience of the shell material to various proteolytic enzymes as well as strong acids and bases even under high temperatures (*14, 15*).

**Fig. 1.**
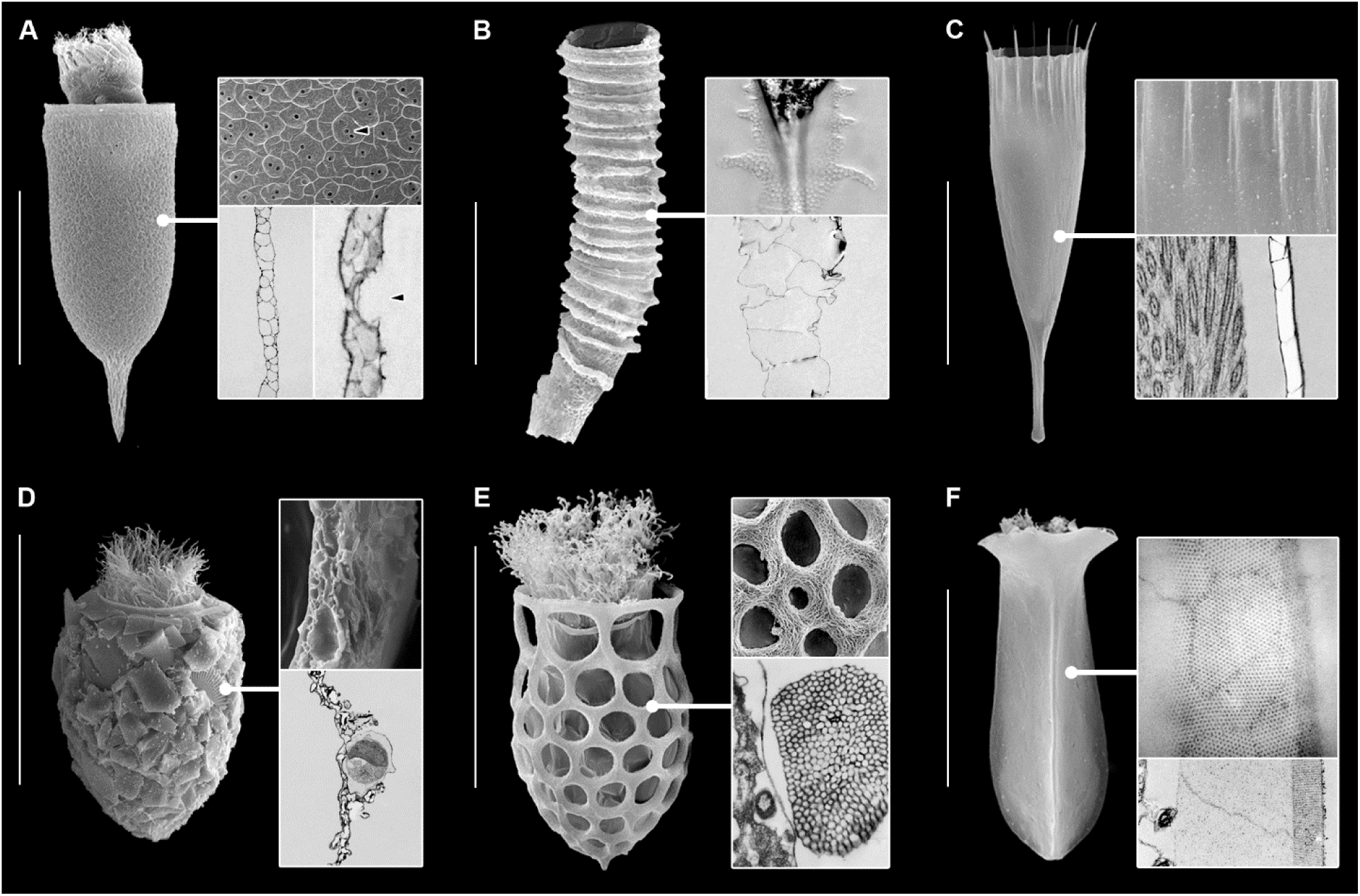
Diversity of shells in tintinnid ciliates. The wall is optically transparent (A–C, E, F) or agglutinated (D) with chambered (A–D), tubular (E), or solid (F) textures. **(A)** *Schmidingerella*, shell with protruding cell (SEM, scanning electron micrograph), detail of outer shell surface (SEM), and two ultrathin sections (TEM, transmission electron micrographs) showing wall texture, surface ridges, and minute pores (arrowheads). **(B)** *Climacocylis*, shell (SEM), detail in the light microscope showing spiraled ridge, and wall section (TEM). **(C)** *Dadayiella*, shell (SEM), detail of anterior outer shell surface (SEM), and wall section (TEM). **(D)** *Stenosemella*, shell (SEM), fracture surface (SEM), and wall section (TEM) with adhered and imbedded foreign particles. **(E)** *Dictyocysta*, shell (SEM), detail of outer surface (SEM), and wall section [TEM (*16*)]. **(F)** *Amphorides*, shell (SEM), tangential section of outer wall layer and transverse section showing the solid wall with a crystalline outer and an amorphous inner layer (TEM). Scale bars are 100 µm (A, B), 50 µm (C–F).

The chemical composition of tintinnid shells has remained a mystery for over two centuries (*14, 15*). The shell-forming material is produced during the cell cycle and accumulates as secretory granules that progressively mature (*17*). Hardly anything is known about shell construction by tintinnid ciliates because the process has rarely been observed or documented, until now. In species forming transparent shells, the process is usually completed within a few minutes after cell division and involves a coordinated interplay of regulated granule secretion, cell movement, and self-assembly of the shell-forming granular material (*17, 18*).

This study identifies the main component of the proteinaceous shells by combining single-cell transcriptomic analyses and mass-spectrometry of the shells from the tintinnid *Schmidingerella* (Fig. 2). We incorporate single-cell transcriptome data from further tintinnid species with structurally diverse shells [this study; (*19*)], including agglutinated forms, and metatranscriptome and genome data from plankton samples collected globally (*20, 21*). Further, we describe potential mechanisms of the material self-assembly and discuss the use of tintinnid ciliates as convenient systems to uncover protein sequence-structure-function relationships that will benefit the design of next-generation biomaterials.

**Fig. 2.**
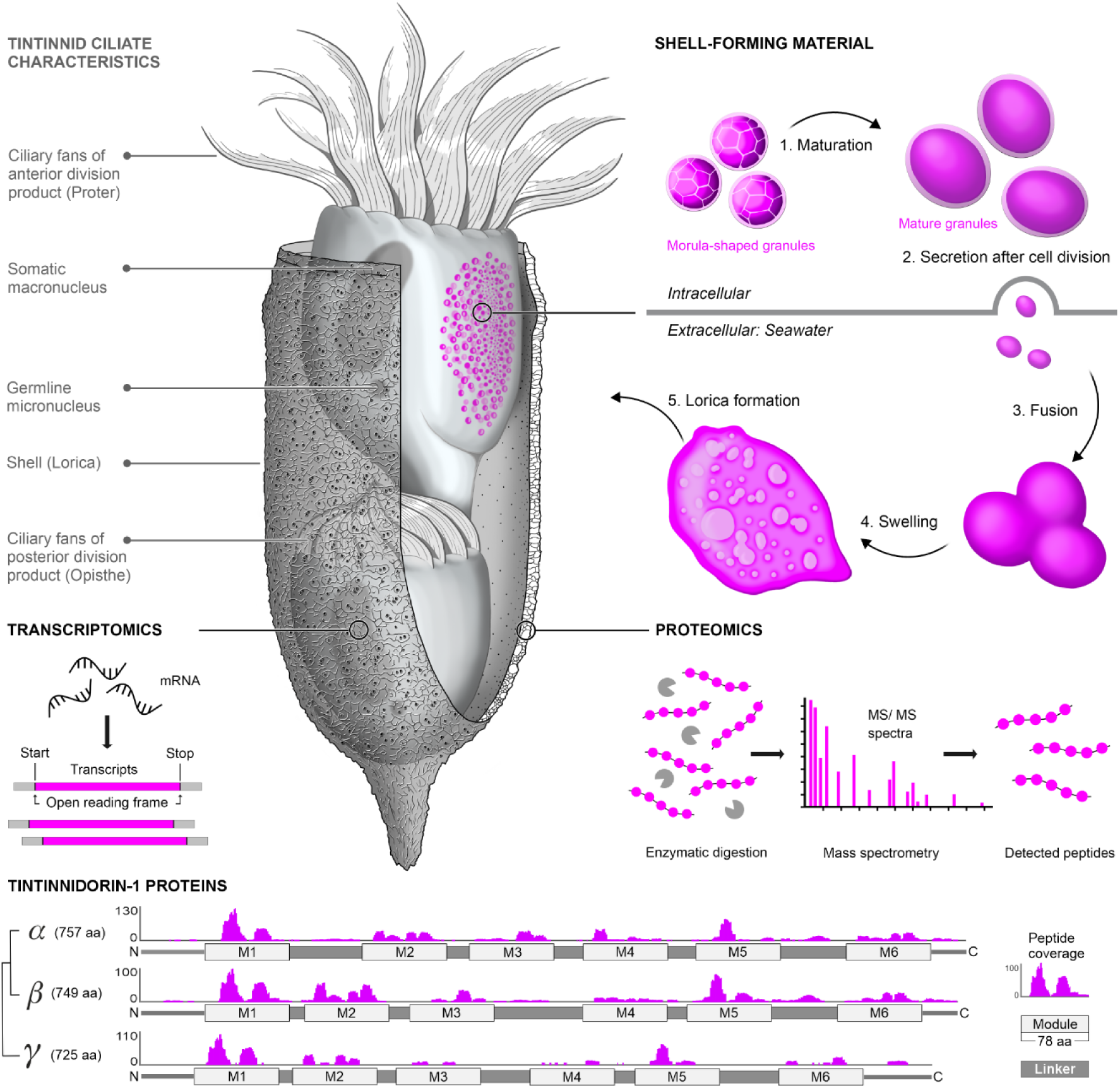
Identification of shell biomaterial proteins in the tintinnid ciliate *Schmidingerella*. Transcriptomic and proteomic analyses on cells and shells congruently identified three variants of a new protein family, namely, Tintinnidorin-1-alpha, beta, and gamma. Gene-sized nanochromosomes encode Tintinnidorin proteins in the macronucleus. During the complex cell division process, messenger RNA (mRNA) is translated into Tintinnidorin proteins that form microscopically visible granules in secretory vesicles. The intracellular granular shell material matures and is secreted by the anterior division product after its separation from the posterior division product, which keeps the parental shell about 200 × 80 µm in size. The rapid self-assembly process involves the fusion and swelling of the extracellular granules, eventually forming the shell of the tintinnid steadily swimming in the water column by means of its apical ciliary fans. The three Tintinnidorin protein variants comprise six modules of identical length alternating with linkers of highly variable lengths. The short chains of amino acids (peptides) identified in the tandem-mass spectrometry data of enzymatically digested *Schmidingerella* shells mainly cover the modules of the three Tintinnidorin protein variants; their quantity is depicted by magenta curves parallel to the protein sequences. Pairwise comparisons reveal a higher sequence similarity between Tintinnidorin-1-alpha and beta as illustrated in the tree (see also fig. S8). aa, amino acids; C, C-terminal region; N, N-terminal region.

### Identification of the shell biomaterial proteins

We obtained single-cell transcriptome data from a monoclonal culture of the tintinnid *Schmidingerella*, including 20 individuals in early, middle, late, and post-division stages of the cell cycle. The transcriptome data from each individual (cells 1–20; table S1) as well as the pooled data from all cells were assembled *de novo* because no complete reference genome or transcriptome exists. The pooled data assembly, termed the “*Schmidingerella* reference transcriptome”, represents our best inference of protein-coding sequences as it comprises the most comprehensive proteome coverage of a tintinnid ciliate’s cell cycle to date. Additionally, to determine their composition, we enzymatically digested about 1,700 of the transparent, champagne flute-shaped shells from the same *Schmidingerella* culture with a custom protocol.

The composition of the shells was assessed after tandem-mass spectrometry. We used a machine learning model (*22*) to identify potential shell peptide sequences from the mass spectra *de novo*, that is, without reference databases of known peptides or any prior information on the protein sequences. We validated the identified shell peptide sequences (Fig. 2) by manual assignment of the mass spectra (data S1 and table S2).

Combining transcriptomic and mass spectrometry data enabled direct identification of the shell proteins by their full-length amino acid sequences. Peptides from the digested shells that were predominant in several independent tandem-mass spectrometry measurements matched various segments of three full-length protein-coding sequences (757, 749, and 725 amino acids in length) of the *Schmidingerella* reference transcriptome (Fig. 2). Two further features of the identified protein sequences provide direct links to the shell-forming material production and secretion in our *Schmidingerella* cells. First, the messenger RNAs (mRNAs) encoding these protein sequences are more highly expressed in the late stages of *Schmidingerella*’s cell division than in early and middle division stages (fig. S1 and table S3). Thus, the gene expression levels mirror previous light microscopical observations of cells from the same culture, in which the largest increase of shell-forming material was quantified in late cell cycle stages (*23*). Secondly, the three protein variants each contain a eukaryotic signal peptide for extracellular transport at the N-terminus determined by protein language model predictions (*24*) (table S4).

We detected homologs of the identified shell proteins in single-cell transcriptomes of several other tintinnid species (table S5). The shell proteins clustered in a single orthogroup that only contained sequences from tintinnid ciliates (fig. S2). In contrast, no further sequence homologs were found in an extensive dataset of 1,000 genomes and transcriptomes from a wide variety of archaea, bacteria, and eukaryotes (*25*) including tintinnid-related planktonic ciliates without shells. Furthermore, structure searches against comprehensive sets of experimentally determined as well as predicted protein structures did not reveal significant structure similarities to known proteins (fig. S3 and table S6). The results strongly suggest that the structural proteins forming the shell biomaterial exclusively evolved in tintinnid ciliates. Here, we term the newly discovered protein family “Tintinnidorin” (supplementary text). Screening the Tara Oceans data in the Ocean Gene Atlas (*26*) and the North Pacific Eukaryotic Gene Catalog (*21*) revealed an extensive phylogenetic diversity of Tintinnidorin proteins greatly expanding our single-cell dataset (Fig. 3 and table S7).

**Fig. 3.**
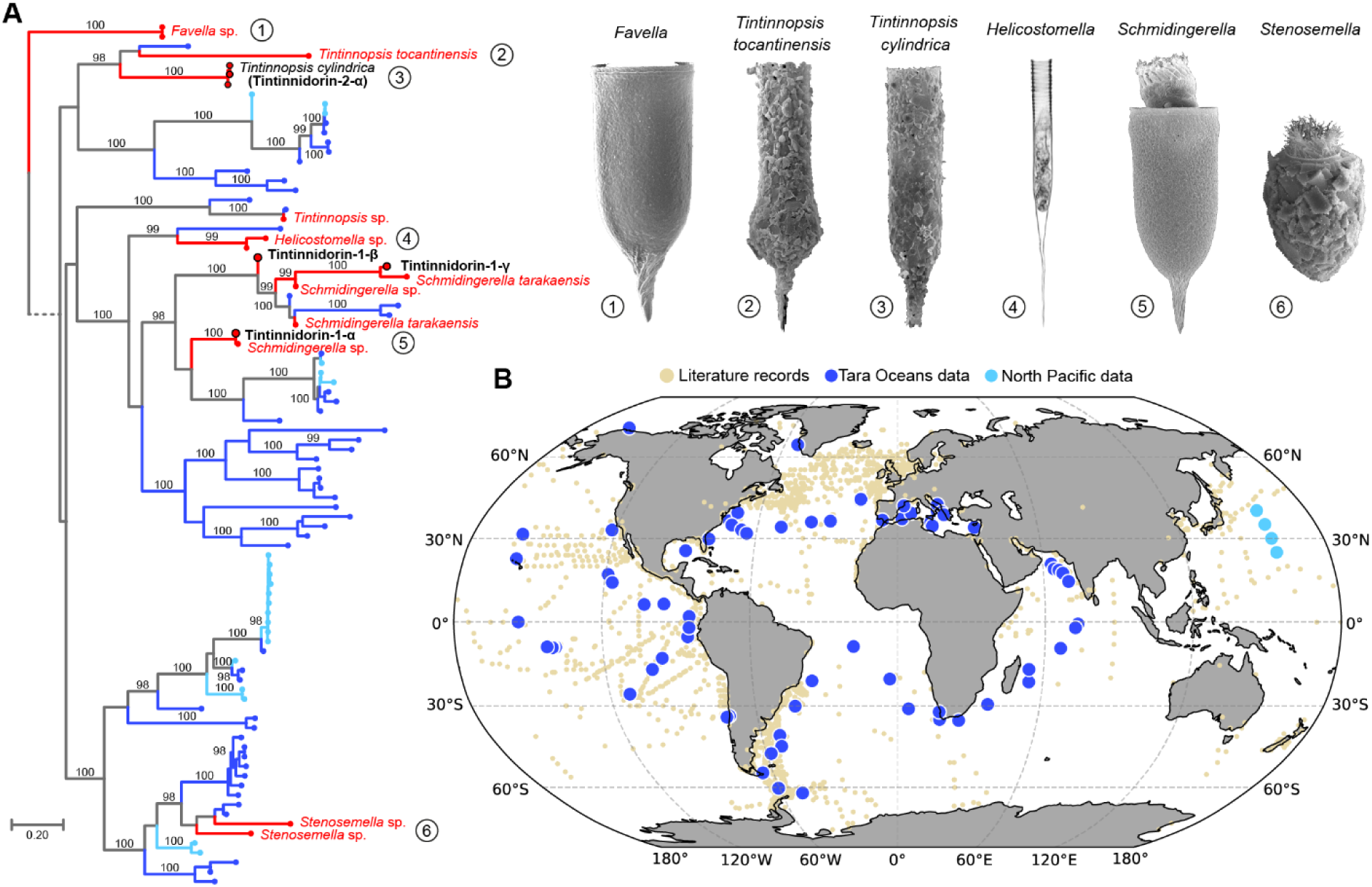
Tintinnidorin proteins extracted from single-cell transcriptomes and discovered in metatranscriptome data from global ocean plankton surveys. **(A)** Phylogeny of Tintinnidorin proteins. Single-cell data annotated with taxon names, light microscopic (4) and scanning electron microscopic images (1–3, 5, and 6) comprise full-length sequences (black) of *Schmidingerella* (Tintinnidorin-1-alpha, beta, and gamma), *Tintinnopsis cylindrica* (Tintinnidorin-2-alpha), and various partial sequences (red). The shells of *Favella*, *Helicostomella*, and *Schmidingerella* are optically translucent, while those of *Tintinnopsis* and *Stenosemella* are agglutinated with mainly mineral particles. The full-length Tintinnidorin sequences from the Tara Oceans data (dark blue) and the North Pacific Eukaryotic Gene Catalog (light blue) are not linked with morphospecies. Scale bar indicates substitution rate. Bootstrap support values (≥ 98) are shown on branches. **(B)** Global distribution of tintinnid ciliates based on the identification of their shells (literature records) and Tintinnidorin proteins.

### Characteristics of Tintinnidorin proteins

Tintinnidorin protein characteristics are the foundation for the multifaceted features and diversity of tintinnid ciliate shells. Our first insights into these characteristics are based on *in silico* analyses that mainly cover amino acid composition, sequence architecture, and predicted protein structures. Physicochemical mechanisms deduced from these analyses elucidate features, such as structural resilience, the capability for wet adhesion, and self-assembly, which follow general principles also associated with diverse terrestrial and aquatic animal biomaterials (*27–30*).

Tintinnidorin protein compositions generally deviate from those of proteins in the Swiss-Prot database (*31*) (Fig. 4A) and the other protein sequences in the *Schmidingerella* reference transcriptome due to high fractions of alanine, glycine, aspartic acid, tyrosine, and tryptophan (Fig. 4A and table S8). While the remaining amino acids are present in varying amounts, cysteine is completely absent in Tintinnidorin-1 proteins of *Schmidingerella* (fig. S4) and rarely existent in those of the other tintinnid species analyzed so far (table S8). The most frequent amino acids in Tintinnidorin proteins are similarly predominant in different components of terrestrial and aquatic animal biomaterials. Alanine and glycine are the most prevalent amino acids in the main structural proteins of terrestrial animal derived silks, such as spider spidroins (*1, 32*) and silkworm fibroins (*33*). Tyrosine and lysine are abundant in proteins of marine animal biomaterials that are related to wet adhesion, such as mussel byssal foot proteins (*34*) and sandcastle worm tube cement (*28*). Aspartic acid mainly occurs in protein components of fibroin-derived silks produced by terrestrial and aquatic animals (*1*).

**Fig. 4.**
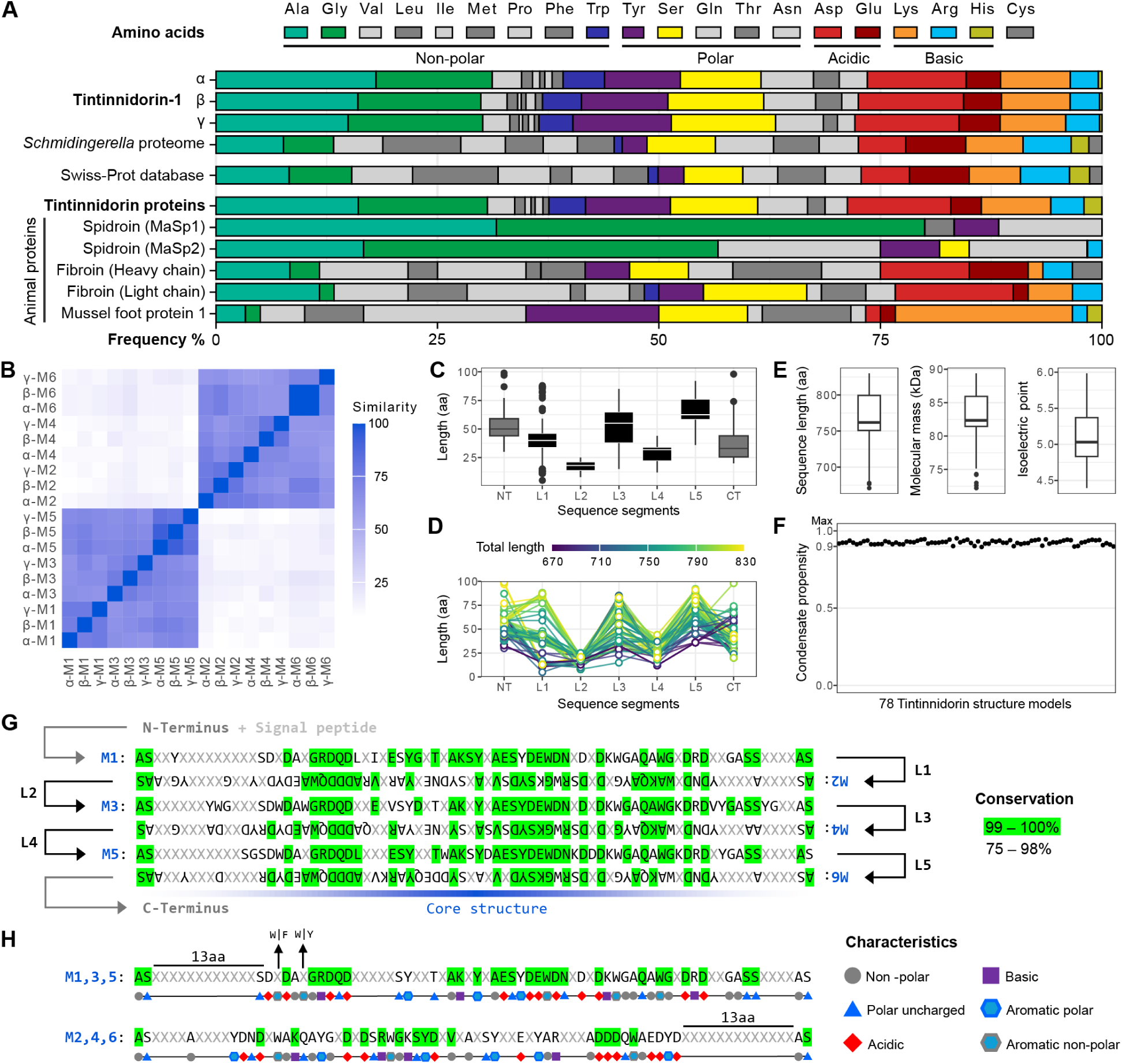
Comparisons of amino acid compositions. **(A)** Amino acid frequencies in Tintinnidorin-1-alpha, beta, and gamma proteins and their average values in the *Schmidingerella* proteome, the Swiss-Prot database, the 78 full-length Tintinnidorin protein sequences, and selected animal biomaterial proteins (spider major ampullate spidroins 1 and 2; silkworm fibroin heavy and light chains; mussel foot protein 1). Especially, high tryptophan frequencies characterize Tintinnidorin proteins. **(B)** Pairwise module comparisons of Tintinnidorin-1-alpha, beta, and gamma from *Schmidingerella* display high sequence similarity values among odd-numbered and even-numbered modules, respectively. **(C, D)** Sequence length distributions of N-terminal (NT), linker (L1–5), and C-terminal segments in the 78 full-length Tintinnidorin sequences. Boxplots (C) as well as datapoints and lines colored according to the total length of each sequence including modules (D) display that odd-numbered linkers comprise generally more amino acids than even-numbered linkers. **(E)** Boxplots of sequence lengths, molecular masses, and isoelectric points of the 78 full-length Tintinnidorin proteins. The low isoelectric points indicate that the proteins have a reduced solubility at low pH values, fostering intracellular protein-protein interactions and stable storage conditions and an extracellular functionality at seawater salinities. **(F)** Condensate (phase separation) propensity values predicted for the 78 full-length Tintinnidorin structure models are close to the maximum value. **(G)** Consensus of the six modules among the 78 full-length Tintinnidorin protein sequences shows a considerable amino acid conservation within and between modules. Conserved amino acids present in ≥ 99% of sequences marked by green, amino acids present in 75–98% of sequences are denoted by their one-letter code, while variable amino acids are indicated by X. Sequential stacking of the sequence segments representing the six modules results in an alignment equivalent to the antiparallel beta-sheet core structure predicted by AlphaFold2. **(H)** Consensus amino acid sequences of odd-and even-numbered modules, respectively, in the 78 full-length Tintinnidorin sequences. Conserved amino acids present in ≥ 99% of sequences are marked by green, amino acids present in 75–98% of sequences are denoted by their one-letter code and annotated with their characteristics, while variable amino acids are indicated by X. α, β, γ, Tintinnidorin-1-alpha, beta, gamma; aa, amino acids; CT, C-terminal segment; L1–5, linker segments 1–5; M1–6, modules 1–6; NT, N-terminal segment.

Tryptophan, one of the rarest amino acids in eukaryotes (*35*), occurs in Tintinnidorin in much higher amounts (3.5–4.7%) than in known proteinaceous biomaterials and most proteins in general (Fig. 4A and table S8). The abundances of the aromatic amino acids tyrosine and tryptophan suggest that the shell biomaterial effectively absorbs UV light. In fact, we observed autofluorescence of *Schmidingerella* shells and transparent shells of other tintinnid species excited with UV light (fig. S5; excitation at 385 nm, emission at about 465 nm). Although excitation and emission wavelengths of these amino acids are typically much shorter (280 nm and 295–400 nm, respectively) (*29, 36*), several processes, e.g., the formation of bonds, oxidation, and structural features of the biomaterial, could result in a red edge excitation/emission shift (*37*).

In terms of sequence architecture, Tintinnidorin proteins consist of six regions with an identical number of amino acids (modules M1–6) alternating with five regions of varying lengths (linkers L1–5) between the N-and C-terminal ends (Fig. 2 - bottom and fig. S6). The linkers are between 5 and 91 amino acids in length (Fig. 4, B and D and table S9), whereas the modules are invariably 78 amino acids long and mainly flanked by pairs of alanine and serine, sometimes aspartic acid or rarely other residues (Fig. 4G). Compositionally, modules and linkers differ significantly in their most abundant amino acids glycine, tryptophan, serine, aspartic acid, glutamic acid, lysine, arginine, and histidine (fig. S7, A and B and table S10–S12). This pattern of modules and linkers distinctly deviates from known structures of animal silk proteins, which mainly comprise low-complexity sequence regions rich in glycine and alanine and arranged in tandem repeats between the compositionally more diverse N- and C-terminal domains (*1*).

Amino acids: A, Ala, alanine; G, Gly, glycine; V, Val, valine; L, Leu, leucine; I, Ile, isoleucine; M, Met, methionine; Pro, proline; F, Phe, phenylalanine; W, Trp, tryptophan; Y, Tyr, tyrosine; S, Ser, serine; Q, Gln, glutamine; T, Thr, threonine; N, Asn, asparagine; D, Asp, aspartic acid; E, Glu, glutamic acid; K, Lys, lysine; R, Arg, arginine; H, His, histidine; Cys, cysteine.

Tintinnidorin modules are not identical in their amino acid sequences but show an alternating pattern with low similarity between pairs of subsequent modules (14 ± 3%) and high similarity between pairs of odd-numbered (70 ± 6%, M1, 3, 5) or even-numbered (69 ± 8%, M2, 4, 6) modules (Fig. 4B and table S13). Within odd-and even-numbered modules, about 35% and 21% of the amino acid positions are conserved across all Tintinnidorin sequences, respectively, and thus represent distinct sequence motifs (Fig. 4, G and H). In contrast, the linkers contain no generally conserved positions, but a mixture of partly repeating amino acid segments mainly composed of glycine, tyrosine, lysine, alanine, and histidine. Additionally, linkers display a significantly higher variance of most amino acids compared to the modules (fig. S7A and table S10). Usually, two to four proline residues are present in each of the N- and C-terminal segments, whereas about two thirds of the Tintinnidorin proteins contain single proline residues in one or more of their linkers. The near absence of linker peptides in the proteomic analyses (Fig. 2 – bottom) suggests that shell digestion by the three proteases either produced linker fragments below the detection limit, or alternatively, that covalent crosslinks impeded proteolytic cleavage, resulting in large and heterogeneous peptides unsuitable for mass spectrometry. Lengths of linkers 1, 3, and 5 vary widely, whereas those of linkers 2 and 4 show less variation in *Schmidingerella* and the other Tintinnidorin sequences. On average, linkers 2 and 4 are also consistently shorter than linkers 1, 3, and 5 (18 and 29 vs. 42, 54, and 66 amino acids) (Fig. 4C and table S9).

The emergence of conserved and variable elements in Tintinnidorin proteins is most likely related to the unusual genome architecture of ciliates, a lineage defined by the presence of both a germline micronucleus and somatic macronucleus in every individual (*38*). One of the three Tintinnidorin variants of *Schmidingerella*, Tintinnidorin-1-gamma, appears to be a canonical paralog as it is divergent from the other two across its full length. In contrast, Tintinnidorin-1-alpha and beta are roughly 30% divergent across two thirds of their lengths; the remaining portions of these two sequences are identical (fig. S8). This pattern is consistent with alternative processing of shared germline sequences (*39, 40*), a process that is associated with elevated rates of protein evolution in ciliates as compared to other eukaryotes (*41, 42*). An alternative explanation/hypothesis is that gene conversion has homogenized the 3’ ends of Tintinnidorin-1-alpha and beta.

Protein structure predictions of the 78 full-length Tintinnidorin sequences by AlphaFold2 (*43*) consistently resulted in monomers with strikingly similar core structures (figs. S9 and S10). The core structure comprises the six modules that form a layer of antiparallel beta**-**sheets connected by the linkers that lack a stable, well-defined three-dimensional structure, indicating that they are intrinsically disordered (Fig. 5A and fig. S11). Similarly, the terminal regions, especially the C-terminus, show a high propensity for disorder (fig. S11). However, the N- and C-terminal segments form two intertwined beta-strands in most monomer models, suggesting these sites to interact with other monomers (Fig. 5A). The predicted protein structures with the module-linker pattern are supported by secondary structure assignments by STRIDE (*44*), disorder predictions by metapredict V2 (*45*), and the AlphaFold2 local confidence scores (Fig. 5, B and C and fig. S6).

**Fig. 5.**
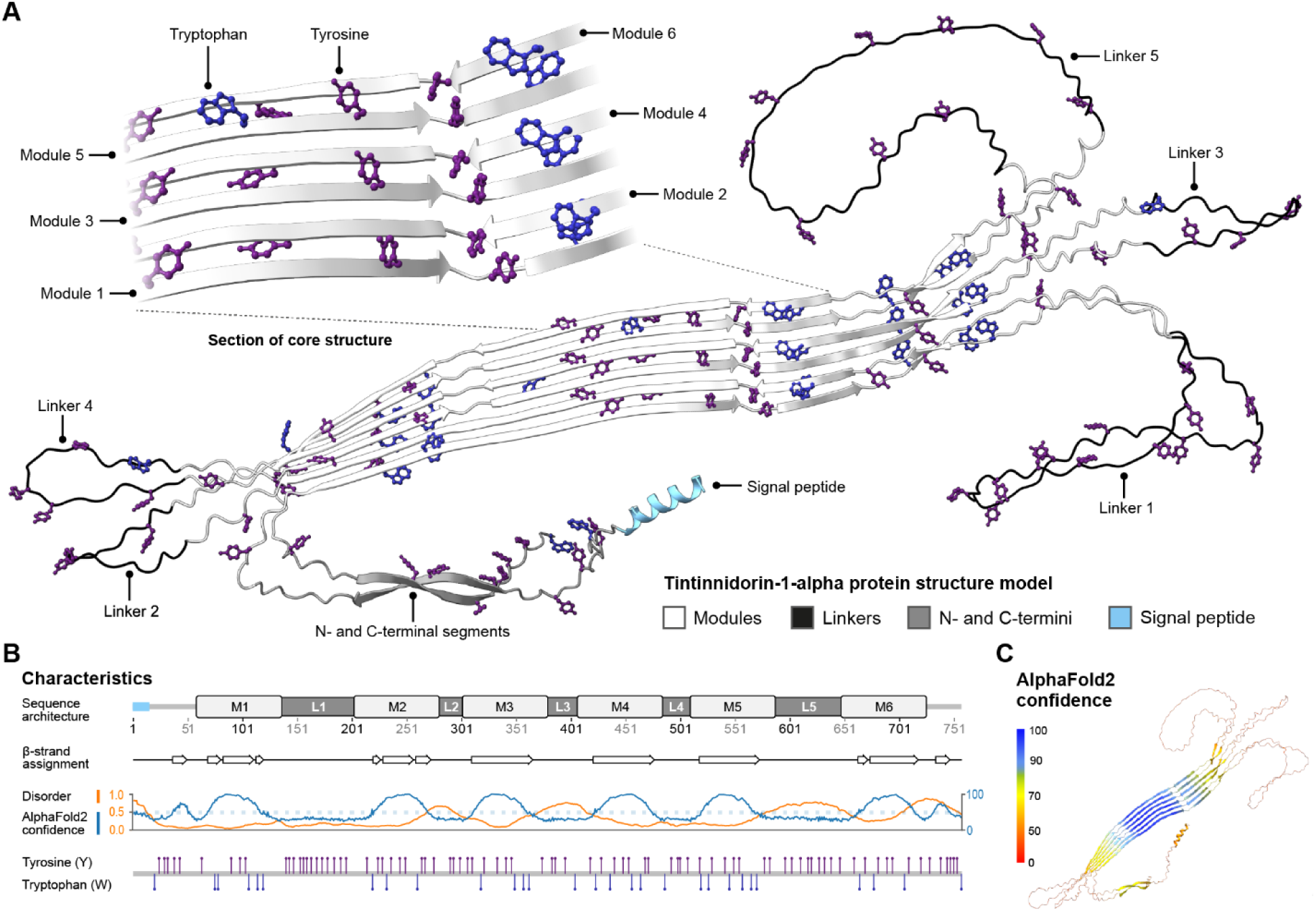
**Predicted protein structure of Tintinnidorin-1-alpha from *Schmidingerella.*** Six antiparallel beta-sheets mainly comprising the modules form the core structure (A) with high support by beta-strand assignments by STRIDE (b) and AlphaFold2 confidence values (B, C; coded by blue). The modules 1–6 (M1–6) are connected by linkers 1–5 of variable length, which display high values of disorder (B) as predicted by metapredict V2, probably causing a high conformational heterogeneity and low model confidences (B, C; coded by orange). The aromatic amino acids tyrosine (Y; purple) and tryptophan (W; blue) are highly abundant and rather homogenously distributed across the protein (A, B). The signal peptide for extracellular transport precedes the N-terminal segment, which forms intertwined beta-strands with the C-terminal segment.

Sequence and structural characteristics of Tintinnidorin proteins are consistent with features displayed by the shell-forming material. Tintinnidorin proteins universally show a high condensate-forming potential, which means that they very likely undergo liquid-liquid phase separation (LLPS; scores ≥ 0.9, on a scale of 0 to 1) (Fig. 4F and table S7). LLPS is strongly associated with storage and self-assembly processes in many protein-based materials of animals and pathogenic amyloids in humans (*1, 27, 46, 47*). The flexibility of Tintinnidorin proteins required for undergoing LLPS and gelation is most likely achieved by the intrinsic disorder of their linkers (fig. S11). Actually, computational simulation experiments suggest that the properties of the intrinsically disordered linker regions, such as length and amino acid-dependent expansion or compaction in solution, can actively influence phase separation and gelation processes (*48*).

Intracellular shell-forming material is stored in secretory vesicles (Fig. 2) that are strongly eosinophilic as revealed by histological stains of *Schmidingerella* (*23*), indicating acidic conditions (low pH ∼4–5) (*49*). Tintinnidorin proteins exhibit low isoelectric points (mean pI = 5.1; Fig. 4E and table S7), resulting in near neutral net charges at low pH, which promotes protein-protein interactions due to reduced solubility and thus stable storage conditions. To concentrate and store functional biomaterial proteins within small volumes, reversible bonds might be formed by stickers (tryptophan, tyrosine, phenylalanine, and histidine residues) that are interspersed between flexible spacers (glycine and alanine residues) according to the sticker and spacer model (*50*). In the Tintinnidorin proteins, the most potent “sticker” amino acids, namely, tryptophan and tyrosine, are abundant and evenly distributed (Fig. 5, A and B and fig. S6), while flexible “spacer” amino acids are mainly present in the linkers (fig. S7B). Thus, interactions between the linkers and the modules under acidic conditions might enable densely packed proteins in the secretory vesicles, avoiding formation of irreversible and harmful assemblies similar to amyloid peptide aggregations (*51*).

Extracellular formation of antiparallel beta-sheets contributes significantly to the mechanical strength and stability of structural proteins in silk biomaterials (*52*). In spiders and silkworms, threads are spun from protein fibers, in which beta-sheet formation is mediated by shear forces, pH, and ion concentration specifically modulated in the silk-producing organs (*27, 53, 54*). Likewise, mussels form proteinaceous byssus threads in a groove of their foot by controlling the microenvironment (*30, 55*). In comparison to these animals, tintinnid ciliates cannot actively control the extracellular physicochemical environment during shell formation as the shell-forming material is rapidly secreted into the surrounding (sea-)water. Nevertheless, family and genus-specific shell characteristics are consistent across a wide range of salt concentrations and water temperatures, suggesting that Tintinnidorin proteins possess features, which maintain their self-assembly capacity.

In many marine prokaryotes, secreted proteins typically have low isoelectric points reflecting adaptations that prevent their denaturation even at seawater salinities (*56*). To our knowledge, Tintinnidorin proteins represent the first example of extracellular eukaryotic proteins that follow this principle to retain their self-assembly function (Fig. 4E). Aromatic residues apparently play an important role for the thermostability of proteins (*57*). In the seawater habitat of *Schmidingerella*, the extracellular conditions are different from the conditions in the secretory vesicles regarding pH (∼8.1), ion composition, and oxidation potential. We observed a fusion and gradual increase in volume (swelling) of the shell-forming material granules (Fig. 2 and movie S1), which are in a viscous gel-like state just after secretion. The hardening of the shell marks the biomaterial’s transition to a solid state, which results in the characteristic chambered texture of the *Schmidingerella* shell wall with its surface ridges and pores (Fig. 1A). Gelation and solidification during shell formation are most likely driven by finely tuned phase transitions including liquid-liquid phase separation (LLPS).

### Global distribution of Tintinnidorin proteins

Tintinnidorin proteins are globally distributed and may serve as a novel marker to systematically assess the biogeography of tintinnid ciliates in future studies. Commonly, the global distribution of tintinnid ciliates has been inferred from their taxonomically relevant shells collected in plankton samples for about 250 years (*58*) (Fig. 3B). In recent decades, molecular data, such as ribosomal marker genes, have significantly enhanced the resolution of tintinnid biodiversity studies through integrative approaches (*12, 59*). We detected Tintinnidorin protein sequences in the Tara Oceans metatranscriptome data acquired from each of the 68 globally scattered sampling sites (*20*) and in the North Pacific Eukaryotic Gene Catalog (*21*) (Fig. 3B and table S14). While the latter comprises samples from a transect extending from the North Pacific Subtropical Gyre to the North Pacific Transition Zone, the Tara Oceans data cover most major oceanic provinces, oceanic and neritic regions, and different water depths (0–1,000 m) (*60*).

These datasets contained 72 full-length Tintinnidorin protein sequences (from start to stop codon) plus many partial sequences (Tara = 370, Pacific = 1,662) that mainly originate from samples taken from the ocean surface layer or at the deep chlorophyll maximum (0–200 m depth).

Taxonomic annotations of the unigenes from the Tara Oceans data to the lowest possible rank (*20*) that we identified as Tintinnidorin sequences show a strong association with ciliates (92%; *n* = 47), and specifically with tintinnid ciliates (45%, *n* = 23; table S7). We note that the approach depends on curated reference databases, which are incomplete, limiting the taxonomic resolution of metatranscriptome sequence annotations. Incomplete recovery of full-length Tintinnidorin transcripts from (meta-)transcriptomes is most likely related to low expression levels, partially degraded mRNA, sequencing bias, and challenges of a *de novo* assembly without reference genomes or transcriptomes, which are common limitations (*61*). Nonetheless, even partial protein sequences could be unequivocally assigned to Tintinnidorin owing to its distinct sequence motifs.

Abundances of tintinnid ciliates vary substantially, from fewer than 100 to over 70,000 individuals per liter, accounting for up to 20% of the total biomass of microzooplankton (20 to 200 µm in size) communities (*62*). One portion of this biomass can be attributed to Tintinnidorin proteins composing the shells, which remain completely intact even after digestion by predators, such as planktonic invertebrates feeding on tintinnid ciliates (*63*). However, the fate of the shells after the decline of a tintinnid population (up to 500,000 sedimenting shells m^-2^ d^-1^) (*64*) and their importance for biogeochemical recycling remain largely unknown. Sparse observations indicate that bacterial degradation may contribute to the breakdown of the tintinnid shell biomaterial during sedimentation or at the bottom of the oceans (*65, 66*).

### Synthesis: Tintinnidorin as a novel model system

Tintinnid ciliates generate an astounding diversity of shells from Tintinnidorin proteins that unite a multitude of remarkable features. Investigating how these features are linked to the distinct characteristics of the shells from molecular to microscopic scales could provide essential insights into protein sequence-structure-function relationships. While general principles of such relationships have been extensively studied in animals (*1, 27, 46*), these complex systems remain inherently challenging, and mimicking biomaterials, such as silk, with comparable mechanical and functional properties has yet to be achieved (*67*).

Tintinnidorin proteins represent a biomaterial with novel architectures that could serve as a convenient model complementing current research. They provide unique insights into the hierarchical self-assembly of proteins into diverse three-dimensional biomaterial layers in water, in contrast to the mostly terrestrial fiber-forming silk biomaterials that are complex mixtures of proteins produced by specialized cells of dedicated spinning organs (*68–71*). How Tintinnidorin proteins interact and finally form a multimeric protein complex in the shell biomaterial will be the aim of future studies. At present, our analyses are restricted to single protein models, as predicting interactions among structural proteins while accounting for changing intracellular and extracellular conditions exceeds current methodological capabilities. In comparison to animals, tintinnid ciliates as single-celled organisms have distinctly shorter generation times (commonly daily cell divisions) and thus rather continuously produce Tintinnidorin proteins. Further advantages are that the cells are directly accessible to staining techniques of subcellular organelles and structures for advanced light and electron microscopic imaging and are potentially suitable targets for genetic engineering.

## Acknowledgements

We thank Michael Gruber (Hieronymus Illustrations) for graphic realization of Fig. 2, Birgit Weißenbacher for discussions about figures and manuscript draft, Taylor R. Sehein for providing Megansett Harbor plankton samples, Ragib Ahsan for guidance on molecular laboratory work, and Xyrus Maurer-Alcalá for input on TIdeS and orthogroup inference.

## Author contributions

Conceptualization: MHG, SA

Funding acquisition and project administration: MHG, SA

Field sampling and culturing: MHG

Shell preparation and enzymatic digestion: MHG, SA

Transcriptomics: MHG

Phylogenomics: MHG, LAK

Mass-spectrometry and data analysis: MHG, CR

Protein analyses and bioinformatics: MHG, MW

Writing – original draft: MHG

Writing – review & editing: MHG, SA, MW, CR, LAK

## Data and materials availability

Raw reads for transcriptomes of our monoclonal *Schmidingerella* (SPMC176) cells generated in this study have been deposited in the NCBI Sequence Read Archive in BioProject PRJNA1363744. Additional single-cell transcriptomes of tintinnid ciliates used in this study are available in BioProject PRJNA1026950 (*19*).

Consensus sequences of the 18S, 5.8S, 28S rRNA genes and the internal transcribed spacer regions (ITS1 and ITS2) of monoclonal *Schmidingerella* specimens (SPMC176) are available in NCBI GenBank under the accession numbers: PX559938 (18S SSU rRNA), PX559939 (ITS1-5.8S-ITS2), and PX644820 (28S LSU rRNA). The mass spectrometry proteomics data have been deposited in the ProteomeXchange Consortium (http://proteomecentral.proteomexchange.org) via the PRIDE partner repository [C] (*114*) with the dataset identifiers PXD070957 and 10.6019/PXD070957. Supplementary Materials (Materials and Methods, tables S1–S14), data (data S1–S3: mass spectra of selected shell peptides, protein and nucleotide sequences of the 78 full-length Tintinnidorin proteins and AlphaFold2 structure models), and Movie S1 are available on Zenodo (10.5281/zenodo.17751168).

## Supplementary Materials

Materials and Methods

Supplementary Text

Figs. S1 to S11

Tables S1 to S14

References (72–114)

Movie S1

Data S1 to S3

