## Supplementary Materials for "A new class of proteinaceous biomaterial with unprecedented properties in unicellular eukaryotes"

**The PDF file includes:**

Materials and Methods
Supplementary Text
Figs. S1 to S11

### Materials and Methods

Sampling and cultivation

Monoclonal cultures of the model tintinnid *Schmidingerella* established with specimens collected from the Rosario Strait, Northeast Pacific were provided by Kelley Bright and Suzanne L. Strom from the Shannon Point Marine Center, Western Washington University, USA, in November 2022. The monoclonal cells investigated in the present study (SPMC 176) were maintained at our laboratory for several months in artificial seawater plus f/2 medium (*72*) at a salinity of 32‰, a pH of 7.8, a temperature of 15 °C, and a light-dark-cycle of 12:12 h. As food items, the dinoflagellate *Heterocapsa triquetra* (CCMP 448) and the haptophyte *Isochrysis galbana* also provided by the Strom Lab were maintained in culture as well. Each well of 6-well plates (CELLSTAR® 6 Well Cell Culture Plates) was filled with 10 ml of filtered ciliate medium and about 5 ml of *H. triquetra* culture and kept for one day in the culture cabinet. Every two or three days, 20–40 *Schmidingerella* cells were transferred into the wells of the prepared plates. Finally, one drop of *I. galbana* culture was added to each well.

Field material sampling and single-cell processing of other tintinnid ciliates (*Favella*, *Tintinnopsis cylindrata*, *T. tocantinensis*, and *Stenosemella*) was performed by colleagues according to methods described in ref. (*19*). Further tintinnid ciliates we collected by a 20-µm-meshed plankton net from surface waters at Avery point, Groton, CT, USA (*Helicostomella*; 41°18'59.2"N 72°03'38.1"W) and Megansett Harbor, North Falmouth, MA, USA (*Metacylis* and *Schmidingerella*; 41°39'22.9"N 70°37'24.8"W).

Fluorescence microscopy

Shells of *Favella*, *Helicostomella*, *Metacylis*, and *Schmidingerella* (Avery Point and Megansett Harbor field samples) were analyzed with a Zeiss Axio Imager M2 microscope equipped with a Zeiss ApoTome.2 device at Smith College in Northampton, MA, USA. Images were captured with a Zeiss Axiocam 503 mono digital camera and the ZEN software v3.8 (Zeiss) at 200× and 400× magnifications and an excitation wavelength of 385 nm (UV light).

Single-cell transcriptomics of *Schmidingerella*

*Isolation of cells*: To reduce contamination by non-target organisms, *Schmidingerella* specimens (*n* = 200) from the monoclonal culture were picked in batches of about 20 cells with a finely drawn glass Pasteur pipette and transferred three times to fresh wells of 6-well culture plates containing exclusively artificial seawater. Cells were either directly processed or starved for 10 min to 24 h. Next, single cells were transferred to a drop of artificial seawater on a disinfected microscope slide to determine the division stage under an Olympus BX53 compound light microscope with differential interference contrast optics at 200–400× magnification without a cover slip.

*Cell classification*: Assignment of specimens to division stages followed the definitions in ref. (*23*). The main distinguishing feature was the shape of the oral primordium (new oral apparatus of the posterior division product): small non-divider without oral primordium or early divider with a very small one (ED); middle divider with crescent-shaped oral primordium (MD); late divider with circular oral primordium parallel to cell surface (LD); and very late divider with oral primordium in distinct division furrow (VLD; Fig. 2). The category ‘postdivider’ (PD) represents posterior division products (opisthe cells) picked just after cell division (cell 9; table S3) and cells apparently missing a shell, which could include anterior division products (proters) that were unable to form a complete shell just after division or that abandoned their shells (cell 1).

*mRNA extraction, amplification, library preparation, and sequencing*: The SMART-Seq® v4 Ultra® Low Input RNA Kit (TaKaRa Bio, USA) was used for single-cell mRNA extraction, transcription, and subsequent cDNA amplification (*73*). Each classified cell was transferred in about 1 µl of artificial seawater to a PCR tube, to which 0.25 µl 10× reaction buffer and 1.15 µl nuclease-free water were added immediately (total volume ~ 2.4 µl). PCR tubes containing the cells were frozen at -80 °C. Twenty out of the 200 picked *Schmidingerella* cells covering each division stage plus postdividers (ED = 2, MD = 10, LD = 3, VLD = 3, PD = 2) were randomly selected and processed, following the manufacturer's protocol, except for using merely one fourth of the recommended reaction volumes. Samples were purified, using the AMPure XP purification system (Beckman Coulter, USA) and quantified with a Qubit 3.0 Fluorometer (Thermo Fisher Scientific, USA). Sequencing libraries were prepared with the Nextera XT DNA Library and Nextera XT Index kits (Illumina, USA). Paired-end sequencing (PE150) was conducted on an Illumina NovaSeq6000 System at Biomarker Technologies GmbH (Münster, Germany).

*Raw read processing and transcriptome assembly*: The quality of raw reads from each *Schmidingerella* single-cell transcriptome was assessed with FastQC v0.12.1 (*74*), and results were compiled with MultiQC v1.15 (*75*). Sequencing adapters and primers from cDNA amplification were removed. Reads were quality-filtered (phred score threshold = 24) and trimmed to a minimum length of 100 base pairs with BBduk from the BBMap package v39.28 (*76*). Next, reads were processed with RiboDetector v0.3.1 (*77*) to identify ribosomal RNA (rRNA) sequences (-e rrna), which were removed and stored in separate FASTQ files (data S4). *De novo* transcriptome assemblies were generated for each cell, using SPAdes v3.15.5 with the rnaSPAdes mode enabled (--rna) and k-mer sizes of 21, 33, 55, and 77 (*78, 79*). In addition, reads from all twenty cells were pooled and assembled, using the same parameters to generate a reference dataset of protein-coding genes across the cell cycle of *Schmidingerella* termed the "reference transcriptome". Raw read processing and assembly metrics are summarized in table S1.

*Ribosomal RNA sequence assembly*: Reads classified as rRNA sequences were assembled, using the same parameters as for the transcriptome assemblies. The resulting contigs were clustered with CD-HIT v4.8.1 (*80, 81*), using the CD-HIT-EST command at 95% similarity (Parameters: -c 0.95 -n 10 -d 0 -M 0 -T 0). The longest contig from each cluster was used to assess identity and completeness against a reference dataset of *Schmidingerella* rRNA sequences with BLASTN v.2.12.0. Contigs with high similarity to the reference sequences were aligned with MAFFT, using the parameters --globalpair, --maxiterate 1000, and --adjustdirectionaccurately. Complete rRNA sequences were then reconstructed by concatenating overlapping contigs for each cell. Consensus sequences of the 18S, 5.8S, 28S rRNA genes and the internal transcribed spacer regions (ITS1 and ITS2) for our investigated monoclonal *Schmidingerella* specimens (SPMC 176) are available in NCBI GenBank under the accession numbers: PX559938 (18S SSU rRNA), PX559939 (ITS1-5.8S-ITS2), PX644820 (28S LSU rRNA). The present paper is the third publication in a series on shell formation in the model tintinnid ciliate *Schmidingerella*. Specimens of the same monoclonal culture (SPMC 176) had been investigated concerning the accumulation of shell-forming material during the cell cycle, its final volume available for shell formation, and a comparison of the maximum amount of intracellular material with the wall volume of the finished shell (*23*). Further, the maturation and secretion of the shell-forming material and the process of shell formation were studied in the same clone (*17*).

*Identification of protein-coding regions*: Open reading frames (ORFs) in the assembled mRNA transcripts were identified with TIdeS v1.1.2 (*82*) (https://github.com/xxmalcala/TIdeS), which employs a machine learning approach. The TIdeS (Transcript Identification and Selection) framework utilizes a classifier trained on features such as coding potential, sequence composition, and ORF length to distinguish true protein-coding sequences from spurious ones. First, a TIdeS model was trained based on the *Schmidingerella* reference transcriptome and a set of reference protein sequences from six diverse eukaryotes prepared with "prep_tides_db.sh". Next, the model was used to infer full-length ORFs in transcript sequences of each assembled transcriptome with default parameters and the ciliate genetic code (-g 6) for the correct translation of codons.

Proteomics of *Schmidingerella* shells

*Preparation of shells*: Empty and intact shells of *Schmidingerella* specimens (SPMC 176) were continuously collected from the culture material, thoroughly washed in dd$H_2O$ to remove salt, and air-dried in 1.5 mL Eppendorf tubes, using a desiccator. Prior to enzymatic digestion, air-dried shells were resuspended in 100 µl molecular grade water and transferred to glass depression slides. Next, shells were washed two or three times, by transferring them through drops of 100 µl molecular grade water on clean depression slides with a finely drawn glass Pasteur pipette. Subsequently, shells were transferred in a minimum volume of water to a glass depression slide with 100 µl of 50 mM TEAB buffer (triethylammonium bicarbonate, pH 8.5, Sigma-Aldrich). Finally, 100 µl of TEAB buffer with the shells were pipetted into a 1.5 mL LoBind® Eppendorf tube. Three tubes with about 1,700 shells in total were prepared for the enzymatic digestion experiments.

*Enzymatic digestion*: Enzymatic digestion experiments were conducted with three proteases (proteinase K, trypsin, and elastase). Samples were incubated at optimal digestion temperatures according to manufacturer's specifications in an Eppendorf Thermomixer C at 1,000 rpm until most shells visibly dissolved in the solution. After digestion, samples were stored at -20 °C until processing for mass spectrometry. In the first digestion experiment, two samples were prepared, using proteinase K (Qiagen Proteinase K solution) and sequencing grade modified trypsin (Promega). The first sample (Sample 1) containing about 690 shells in 100 µl TEAB buffer was incubated with 3 µl proteinase K (0.3 mg/ml) at 57 °C for 7 h and subsequently with 1 µl trypsin (1 mg/ml) at 37 °C overnight. The second sample (Sample 2) containing about 490 shells in 100 µl TEAB buffer was incubated solely with 2 µl proteinase K (0.3 mg/ml) at 57 °C for 4 h. In the second digestion experiment (Sample 3), about 500 shells were incubated with 2 µl elastase (1 mg/ml; Promega) in 100 µl TEAB buffer at 37 °C for 18 h. Control samples containing identical amounts of TEAB buffer and proteases without shells were incubated alongside the shell digestion samples.

*Tandem-mass spectrometry*: The shell protein digests (50 µL of 100 µL) were acidified, using 2.50% (v/v) aqueous trifluoroacetic acid (Sigma-Aldrich) to a final concentration of 0.20% (v/v). Subsequently, the digests were purified, using Pierce™ C18 Tips (Thermo Fisher Scientific) according to the manufacturer's protocol. The purified peptides were dried at 45 °C and 850 rpm in a Savant SpeedVac™ SPD210 Vacuum Concentrator (Thermo Fisher Scientific) equipped with a VLP120 vacuum pump and a RVT5105 refrigerated vapor trap (Thermo Fisher Scientific). The dried samples were resuspended in 20 µL of 1% (v/v) acetonitrile (ACN; Thermo Fisher Scientific) and 0.1% (v/v) formic acid (FA; ≥ 98%; Sigma-Aldrich) in water, which corresponds to the starting conditions in the ultra-high-performance liquid chromatography (UHPLC) method.

To elucidate their respective sequences, the peptides generated by the various enzymatic digestions of the *Schmidingerella* shells (Samples 1–3) were separated employing reversed-phase UHPLC on a C18-based nano-HPLC column and subsequently analyzed by tandem-mass spectrometry, employing a hybrid quadrupole-orbitrap mass spectrometer (Thermo Fisher Scientific) in data-dependent acquisition mode.

*Detailed sample preparation*: Chromatographic separation of 2 µL sample was carried out, employing reversed phase HPLC on a Vanquish Neo™ UHPLC system (Thermo Fisher Scientific), using an Aurora Ultimate TS C18 column (250 × 0.075 mm i.d.; dp 1.7 µm) from IonOpticks. For the separation, 0.1% aqueous formic acid (solvent A) and 0.1% formic acid in acetonitrile (solvent B) were pumped at a flow rate of 300 nL per minute in the following order: 1% B for 5 min, a linear gradient from 1–5% B in 5 min, a second linear gradient from 5–25% B in 75 min, and a third linear gradient from 25–35% B in 5 min. This was followed by flushing at 80% B for 5 min and column re-equilibration at 1% B for 15 min. The column temperature was kept constant at 50 °C, the autosampler was kept at 7 °C.

The UHPLC system was hyphenated to a Q Exactive™ Hybrid Quadrupole-Orbitrap™ mass spectrometer via a Nanospray Flex™ ion source (both from Thermo Fisher Scientific). The spray voltage was set to 1.5 kV, S-lens RF level to 60.0, and capillary temperature to 250 °C. To increase tandem-mass spectra sampling depth and maximize the coverage of possible precursor ions and subsequently peptides, each sample was analyzed in triplicate. First, each scan cycle consisted of a full scan at a scan range of *m/z* 350–1,200 at a resolution setting of 70,000 at *m/z* 200, followed by 15 data-dependent higher-energy collisional dissociation (HCD) scans, using an isolation window of 1.0 *m/z* at 29% normalized collision energy at a resolution setting of 17,500 at *m/z* 200. Second, each scan cycle consisted of a full scan at a scan range of *m/z* 600–1,800 at a resolution setting of 70,000 at *m/z* 200, followed by 15 data-dependent higher-energy collisional dissociation scans, using an isolation window of 1.0 *m/z* at 25% normalized collision energy at a resolution setting of 17,500 at *m/z* 200. Third, each scan cycle consisted of a full scan at a scan range of *m/z* 1,000–2,200 at a resolution setting of 70,000 at *m/z* 200, followed by 15 data-dependent higher-energy collisional dissociation scans, using an isolation window of 1.0 *m/z* at 29% normalized collision energy at a resolution setting of 17,500 at *m/z* 200. The automatic gain control (AGC) target was set to 1e6 charges with a maximum injection time of 100 ms for the full scan and 5e5 charges and 60 ms for the HCD scans in all three methods. Already isolated precursor ions were dynamically excluded for fragmentation for 10 s. Data acquisition was conducted, using Thermo Scientific™ Chromeleon™ 7.2 CDS (Thermo Fisher Scientific).

Identification and analyses of Tintinnidorin proteins

*Combining mass spectrometry and transcriptome data*: Raw mass spectra data files were converted to the mzML format with ThermoRawFileParser v1.4.4 (*83*) (https://github.com/CompOmics/ThermoRawFileParser) and subsequently processed with Casanovo v4.2.0 (*22*) (https://github.com/Noble-Lab/casanovo), using default weights (*84*) and settings. Casanovo is a machine learning model for *de novo* peptide sequencing that does not rely on reference sequences. Peptide sequences identified from the mass spectra were then matched to protein sequences in the *Schmidingerella* reference proteome. Across the entire reference proteome, only three full-length protein sequences (Tintinnidorin-1-alpha, beta, and gamma) received high numbers of peptide matches and were therefore selected for subsequent analyses.

To quantify the peptide coverage of the identified protein sequences and proteases, the Casanovo output was further processed with the Stitch software v1.5.0 (*85*) (https://github.com/snijderlab/stitch), using Tintinnidorin-1-alpha, beta, and gamma, proteinase K (Uniprot entry: P06873), trypsin (Uniprot entry: P00761), and elastase (Uniprot entry: P00772) as templates. To assess the confidence of the *de novo* sequencing results, selected peptide sequences identified by Casanovo were additionally evaluated by manual inspection of tandem-mass spectra with a fragment match tolerance of 15 ppm of predicted and measured fragments, using the proteomics data viewer PDV (*86*). Furthermore, a mirror plot of the tandem-mass spectra was predicted on the respective peptide sequences based on AlphaPept (*87*), using settings matching the measurements for each spectrum, such as precursor charge and fragmentation energy, allowing comparison between experimental spectra and corresponding *in silico* predictions.

*Cellular localization*: Subcellular localization and sorting signals of Tintinnidorin proteins were predicted with DeepLoc v2.1 (*24*) (https://services.healthtech.dtu.dk/services/DeepLoc-2.1; accessed on 04/17/2024; option "High quality"). The transformer-based protein language model differentiated between membrane-associated and soluble proteins. Protein sequences with predicted sorting signal peptides were additionally analyzed with SignalP v6.0 (*88*) (https://services.healthtech.dtu.dk/services/SignalP-6.0/; accessed on 04/24/2024; option "Eukarya", only predicts Sec/SPI "standard" secretory signal peptides).

*Transcript level expression*: Transcript abundances of Tintinnidorin-1-alpha, beta, and gamma for each *Schmidingerella* cell were quantified by transcripts per million (TPM) values estimated with Salmon v1.10.0 (*89*). TPM values were determined from filtered and quality trimmed paired-end reads (FPE and RPE) with the most conservative settings (--validateMappings, --gcBias, --seqBias, --posBias, --mimicStrictBT2) to remove reads with indels and to account for compositional and coverage biases. Transcript level expression values per cell were extracted from Salmon's "quant.sf" files with tximport (*90*) in RStudio (2023.12.1+402) (*91*) and plotted with pheatmap v1.0.12 (https://github.com/raivokolde/pheatmap).

*Orthogroup inference and homology search*: Orthogroup inference of tintinnid ciliate protein-coding genes (20 *Schmidingerella* cells, the *Schmidingerella* reference transcriptome, and 46 single-cell transcriptomes of other tintinnid ciliates) included 232 genomes and transcriptomes from a wide variety of bacteria, archaea, and eukaryotes (table S5). Coding sequence files (CDS) of genomes were translated to amino acid sequences with the appropriate genetic code, using a custom script (https://github.com/Katzlab/EukPhylo/blob/main/PTL1/Genomes/Scripts/3_GCodeTranslate.py). Open reading frames of protein-coding sequences in transcriptomes mainly comprising ciliates were determined with TIdeS, using the model based on the *Schmidingerella* reference transcriptome. Partial Tintinnidorin sequences that did not comprise the entire ORF were identified by sequence similarity searches against full-length Tintinnidorin sequences (CD-HIT and BLAST) and manually aligned to determine the correct reading frame. Subsequently, partial sequences were translated to amino acid sequences and added to the respective single-cell transcriptome assemblies. Orthogroups and hierarchical orthogroups were identified with OrthoFinder v2.5.5, using default settings (*92*) (https://github.com/davidemms/OrthoFinder). In short, OrthoFinder performs an all-versus-all DIAMOND search to cluster genes into orthogroups, from which gene trees are computed. Next, gene trees are concatenated to infer a rooted species tree. Finally, hierarchical orthogroups are defined by reconciling gene trees with the species tree, identifying the descendant genes of each ancestral gene at successive internal nodes.

More extensive homology searches were conducted with EukPhylo v1.0 (*25*). Lineage-specific orthogroups identified with OrthoFinder were used to generate a database of tintinnid ciliate genes including Tintinnidorin proteins with DIAMOND v2.1.8 (*93*). The database was used to perform a systematic and sensitive search for potential homologs across a curated dataset of 1,000 diverse species of bacteria, archaea, and eukaryotes, utilizing the EukPhylo pipeline. The curated dataset comprised standardized genome and transcriptome assemblies (*94*) (https://doi.org/10.6084/m9.figshare.25336129.v2), some of which were generated from raw data in the sequence read archive (SRA) not otherwise available in GenBank.

BLAST searches of Tintinnidorin-1 sequences against the NCBI GenBank (last accessed October 2025) yielded hits to four sequences of hypothetical proteins with 54–66% sequence identity and partial coverage from a metagenomic dataset annotated by the NCBI Prokaryotic Genome Annotation Pipeline (Accessions: MCP4570280, MCP4287431, MCP4556353, and WP_288100621). The dataset originates from a study that investigated foraminifera and their associated microbiomes in marine sediments (*95*). We assume, the sequenced samples must have contained DNA/RNA from tintinnid ciliate cells or their resting stages (cysts).

Additionally, we identified full-length Tintinnidorin sequences highly similar to those of *Schmidingerella* in the transcriptome data of the marine planktonic ciliate *Strombidinopsis acuminata* (Biosample accession: SAMN02740368). *Strombidinopsis* is closely related to tintinnid ciliates but does not form a shell. The species was cultured and sequenced along with tintinnid ciliate species, including *Schmidingerella* specimens, in the MMETSP project (Bioproject accession: PRJNA248394), and its transcriptome assembly shows high contamination with non-target sequences as evident from codon usage bias analyses in a recent phylogenomic study of planktonic ciliates (*19*). In fact, the codon usage bias plot for *S. acuminata* (*Strombidinopsis* sp. MMETSP0126) displays many transcripts [grey dots; fig. S5 in ref. (*19*)] that match tintinnid ciliate transcripts regarding GC content at the third codon position (GC3) in contrast to transcripts attributed to *S. acuminata* (red dots). Consequently, we excluded this dataset from our analysis.

*Public sequence repository search*: Two public repositories were screened for homologs to Tintinnidorin sequences. The Ocean Gene Atlas (*26, 96*) (https://tara-oceans.mio.osupytheas.fr/ocean-gene-atlas/) was searched, using Tintinnidorin nucleotide sequences as queries against the MATOUv1+T dataset (*20*) with default settings. This dataset contains 116 million unigenes that were obtained from the Tara Oceans expedition (*97*) and assembled with reads from metatranscriptomes clustered at 95% identity. Potential sequence homologs were downloaded from the web service for further processing (last accessed on 05/12/2025). Additional sequences were acquired from the North Pacific Eukaryotic Gene Catalog (NPEGC) (*21*) repository v0.92 (https://zenodo.org/records/13826820). Potential sequence homologs from both datasets were processed in AliView v1.28 (*98*) to determine the correct open reading frame, using Tintinnidorin-1-alpha, beta, and gamma sequences as reference. Only sequences containing the complete reading frame ($n$ = 72), i.e., from start to stop codon, were retained for global distribution mapping and phylogenetic analyses. The locations of metatranscriptome samples, in which reads of full-length Tintinnidorin sequences were detected (table S14), and literature records of tintinnid ciliate occurrences (according to ref. (*58*), pers.

commun.) were plotted on a world map with Matplotlib v3.10 (*99*) and the package Cartopy v0.24.1 (https://doi.org/10.5281/zenodo.1182735).

*Phylogenetic analyses*: Full-length Tintinnidorin sequences detected in the two public repositories (*n* = 72) plus Tintinnidorin-1-alpha, beta, gamma (*Schmidingerella*) and Tintinnidorin-2 (*Tintinnopsis cylindrica*) sequences from the single-cell transcriptomes (*n* = 6) were aligned with MAFFT v7.526 (*100*), using the Smith-Waterman algorithm (--localpair) with 1,000 iterations (--maxiterate 1000). Subsequently, partial sequences (*n* = 12) from further single-cell transcriptomes (*Favella*, *Helicostomella*, other *Schmidingerella*, *Stenosemella*, and *Tintinnopsis* species) were added to the alignment of full-length sequences, using the "--addfragments" command. Maximum likelihood tree inference was performed in IQ-TREE v3.0.1 (*101*) under the variable time substitution model (*102*), a proportion of invariable sites, and a FreeRate model with five categories for rate heterogeneity (-m VT+F+I+R5). Branch support was assessed, using ultrafast bootstraps (-b 1000) and SH-like approximate likelihood ratio test (-alrt 1000) with 1,000 replicates. The tree was plotted and graphically edited with TreeViewer v2.2.0 (*103*).

Tintinnidorin protein characteristics and structure predictions

*Sequence architecture and amino acid composition*: The structuring of Tintinnidorin protein sequences into module and linker regions was based on manual assessment of multiple sequence alignments. In the alignments, module regions consistently formed sequence blocks of 78 amino acids, mostly starting and ending with pairs of alanine and serine (sometimes alanine and aspartic acid, or rarely other residues). Regions between the modules were subsequently defined as linker regions. Amino acid compositions were calculated as fractions based on the overall protein sequence and separately on the module and linker regions. Significant differences of mean fractions and variances were tested with the Wilcoxon signed-rank test as implemented in R and the Levene's test (https://doi.org/10.32614/CRAN.package.car), respectively. Amino acid conservation for each position in the modules was determined by assessing the consensus annotation of the aligned full-length Tintinnidorin protein sequences in Jalview v2.11.5.0 (*104*).

*Structure prediction with AlphaFold2*: A local installation of AlphaFold2 (*43*) was used to predict the three-dimensional structures of full-length Tintinnidorin sequences detected in the two public repositories (*n* = 72) plus Tintinnidorin-1-alpha, beta, gamma (*Schmidingerella*), and Tintinnidorin-2 (*Tintinnopsis cylindrica*) sequences from the single-cell transcriptomes (*n* = 6). From the five models generated by AlphaFold2 for each sequence, respective top-ranked models were used for subsequent analyses.

*Structure similarity search*: Since protein structure is generally more conserved than protein sequence (*105*), structure-based searches were performed in addition to the sequence-based searches for identifying proteins with structures similar to the 78 Tintinnidorin models (queries). We used Foldseek (*106*) easy-search (version 427df8a6b5d0ef78bee0f98cd3e6faaca18f172d; --exhaustive-search 1) to search against all experimentally determined structures available from the Protein Data Bank (PDB; https://RCSB.org) (*107*) as of 11/05/2025 (1,051,375 protein chain targets) and against AlphaFold2 models for a representative set of all canonical sequences in the UniProt database (*108*) as of 07/17/2024 (53,665,860 protein chain targets). Furthermore, we used TopMatch (*109*) for structure searches against the abovementioned set of experimentally determined structures and against all AlphaFold2 models for proteins in the UniProt/Swiss-Prot database as of 10/19/2022 (542,378 protein chain targets). TopMatch was also used to re-align the top-ranked hits obtained by Foldseek to obtain consistent structure similarity scores across all structure searches. The extent of structure similarity was then quantified by TopMatch's structure similarity score (*109*). Model analysis and molecular graphics were performed with UCSF ChimeraX (*110*).

*Prediction of protein features*: Secondary structure elements in the AlphaFold2 models were determined by STRIDE (*44*). Disordered protein regions were predicted with metapredict V2 (*45*). Propensities for condensate formation were predicted with PICNIC (*111*). Molecular masses and isoelectric points were calculated with ProtParam (*112*). Schematic views of Tintinnidorin protein features were generated with the pgfmolbio package (http://www.ctan.org/pkg/pgfmolbio; accessed on 11/20/2025) and TeXshade (*113*).

### Supplementary Text

Nomenclature of newly discovered Tintinnidorin proteins.

*Etymology*: Tintinni-dorin. The protein is unique to the monophyletic tintinnid ciliates. The stem of the taxon's name is combined with the word "dorin", which has roots in different languages (Greek: dôron, neuter, noun). Its core meaning revolves around the concept of a gift handed down through generations.

Full-length Tintinnidorin proteins from single-cell transcriptomes of identified specimens were named according to a hierarchical naming system. Tintinnidorin and a numerical identifier provide a unique label, independent of the taxonomic nomenclature, and Greek letters indicate the variant within that group. Consequently, we named the shell proteins discovered first in *Schmidingerella* Tintinnidorin-1-alpha, beta, and gamma and those of *Tintinnopsis cylindrica* Tintinnidorin-2-alpha.

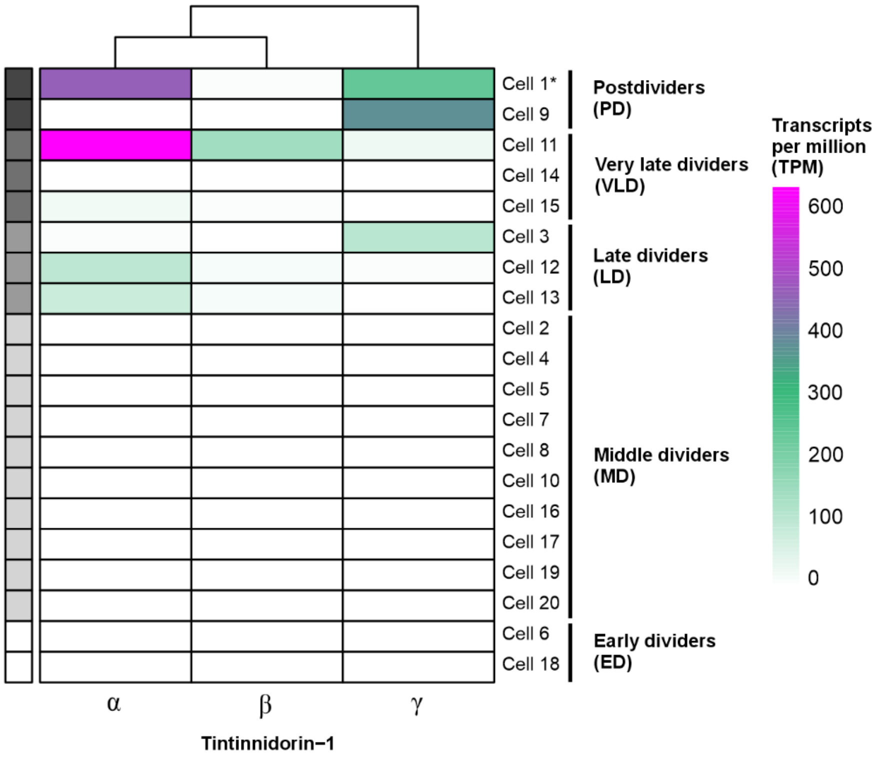


**Fig. S1. Relative gene expression of Tintinnidorin-1-alpha, beta, and gamma proteins during the cell cycle of monoclonal *Schmidingerella* specimens.** The relative gene expression given in transcripts per million is highest in most late dividers (LD), very late dividers (VLD), and postdividers (PD), while earlier stages, namely, early dividers (ED) and middle dividers (MD), generally do not express the genes (table S3). High expression levels of Tintinnidorin are found in a specimen (asterisk; cell 1) that had no shell at the time of sampling but produced proteins potentially for a replacement shell. In another specimen, cell division was observed, and the posterior division product (cell 9) was picked. The three variants are not equally expressed, i.e., transcripts of Tintinnidorin-1-alpha and gamma are more abundant than those of Tintinnidorin-1-beta. Top branching illustrates the sequence similarity of the Tintinnidorin-1 proteins.

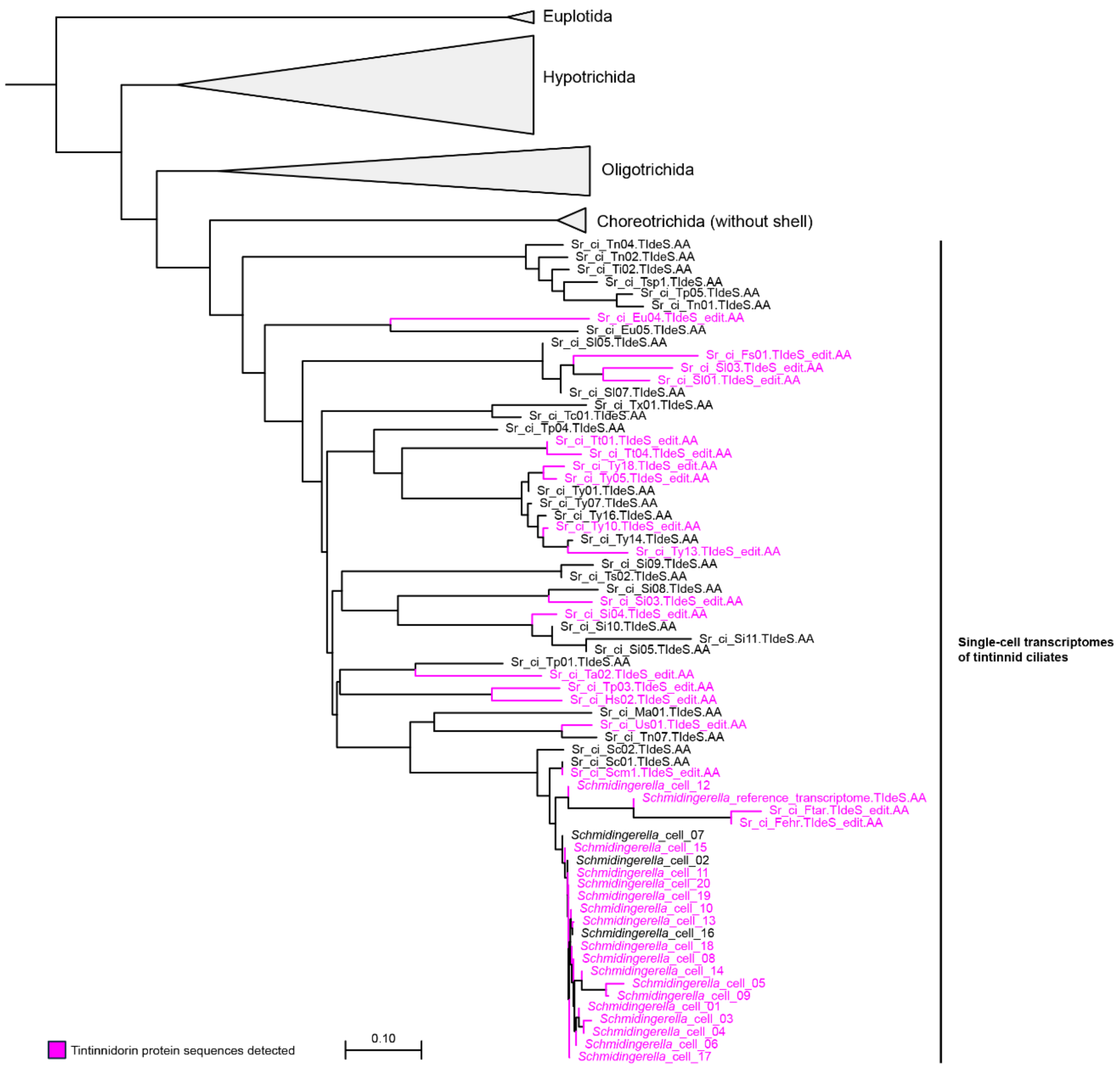


**Fig. S2. Species tree computed by Orthofinder2 based on 298 genomes and transcriptomes** (table S5). Only the tree section including the closest relatives to tintinnid ciliates is displayed. Partial and complete Tintinnidorin protein sequences were exclusively detected in single-cell transcriptomes of tintinnid ciliates (magenta) and assigned to a single hierarchical orthogroup. The tree topology regarding Euplotida, Hypotrichida, Oligotrichida, and Choreotrichida perfectly match phylogenies based on the analyses of nuclear marker genes. Scale bar represents the average number of substitutions per site for a unit branch length.

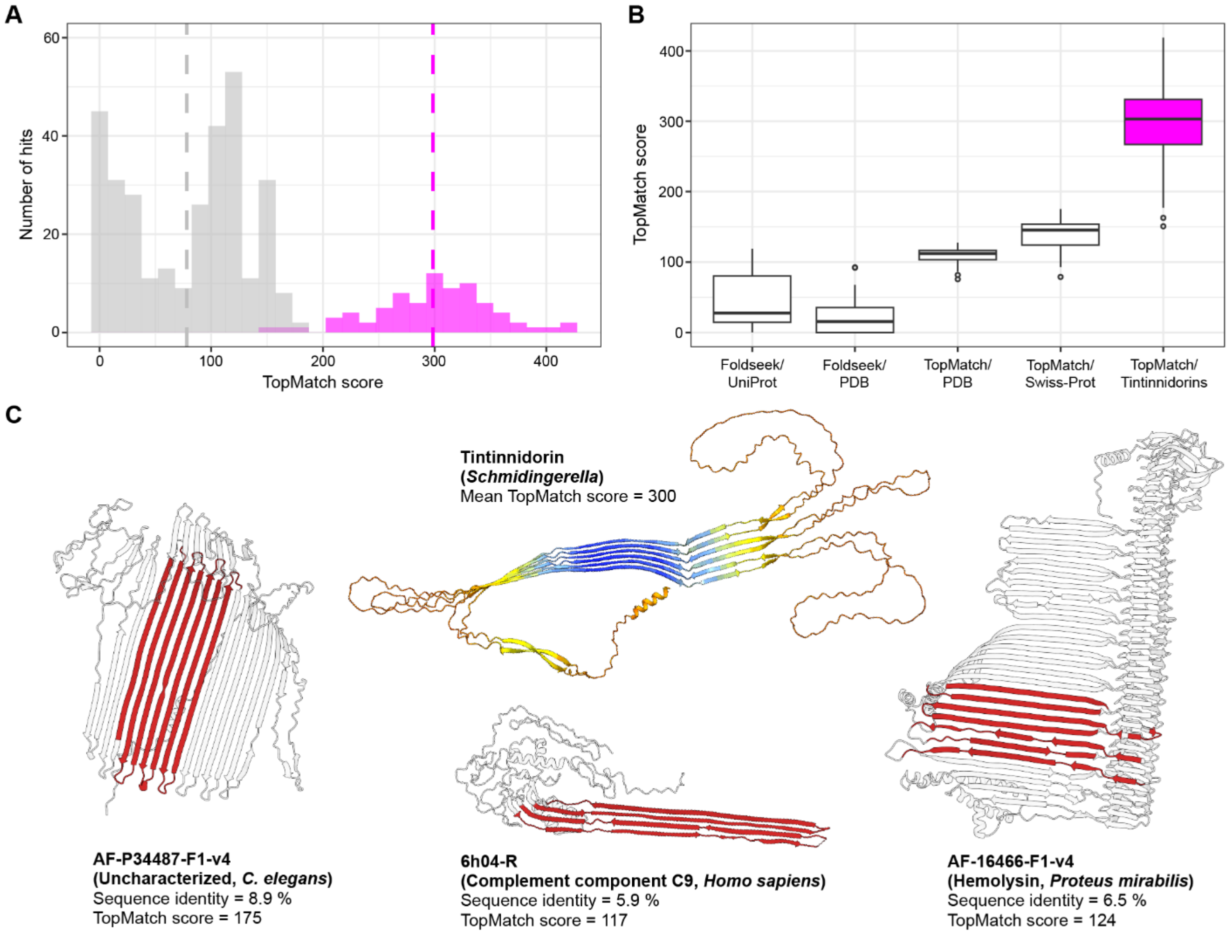


**Fig. S3. Structure similarities of the 78 full-length Tintinnidorin proteins. (A)** Distribution of structure similarity scores as calculated by TopMatch to the best matching targets in exhaustive structure searches against Protein Data Bank (PDB) and UniProt (grey) and in all-against-all structure comparisons of the 78 Tintinnidorin tintinnid shell proteins (magenta). Dashed lines indicate the mean scores for the respective comparisons. While the overall best match of a Tintinnidorin structure to a PDB/UniProt protein structure gets a similarity score of about 175 (table S6), the extent of structure similarity between pairs of Tintinnidorin proteins is, on average, almost twice as high and mainly caused by the structural equivalence of the modules' beta-sheets, with score variance mainly reflecting loop disorder in the linkers. **(B)** Distribution of TopMatch structure similarity scores categorized by search method and target database. **(C)** Structure models of the three best matches to Tintinnidorin proteins based on the aligned segments shaded in red. The low sequence identities and TopMatch scores show that Tintinnidorin proteins have no structural homologs in any protein database.

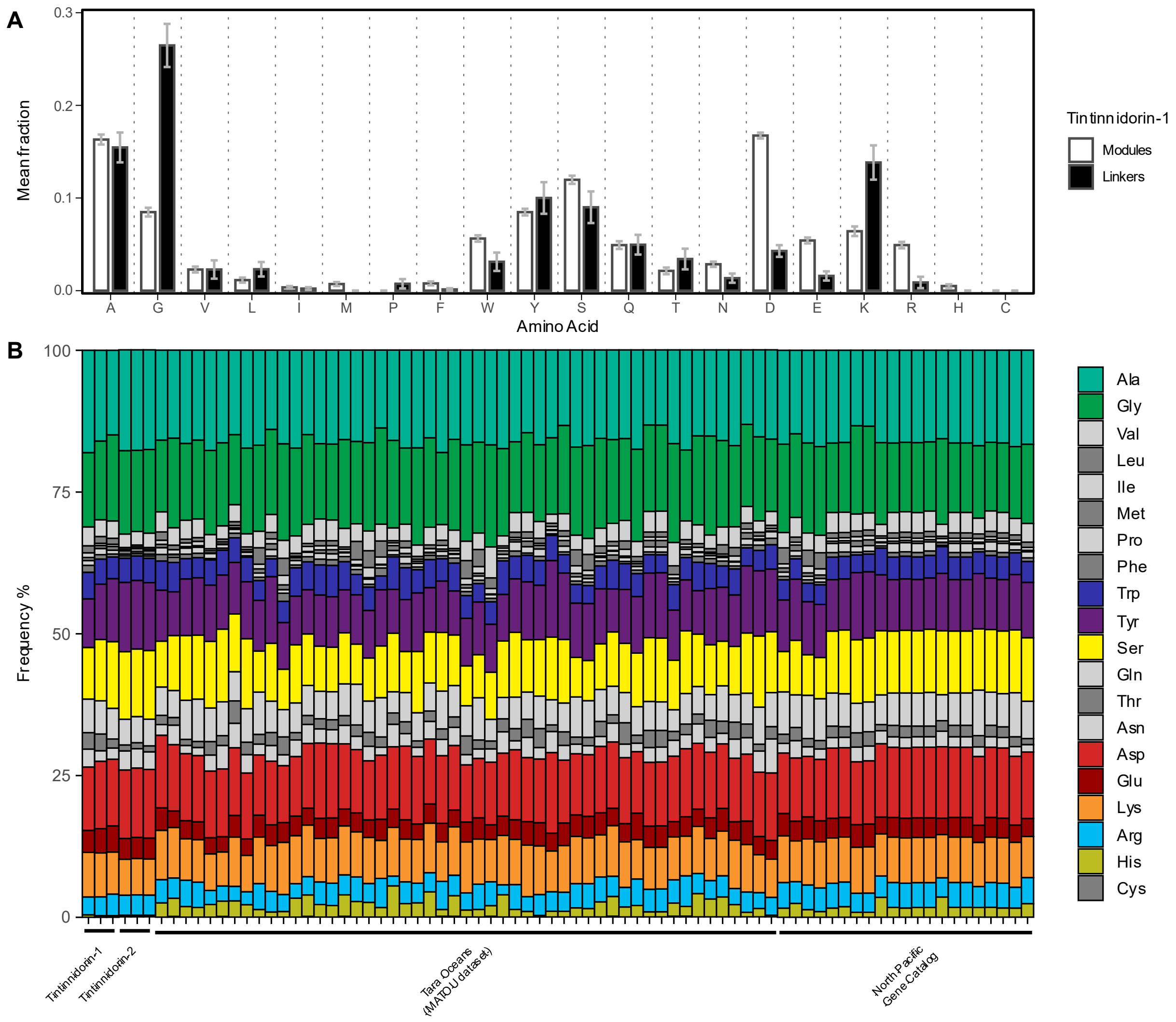


**Fig. S4. Comparisons of amino acid compositions. (A)** Comparisons of the amino acids' mean fractions between modules and linkers in the Tintinnidorin-1-alpha, beta, and gamma proteins of *Schmidingerella*. Error bars = standard error. **(B)** Amino acid frequencies in the 78 full-length Tintinnidorin proteins, including Tintinnidorin-1 of *Schmidingerella*, Tintinnidorin-2 of *Tintinnopsis cylindrica*, and the sequences discovered in the Tara Oceans database and North Pacific Eukaryotic Gene Catalog, demonstrate the compositional similarity of the sequences. A, Ala, alanine; G, Gly, glycine; V, Val, valine; L, Leu, leucine; I, Ile, isoleucine; M, Met, methionine; P, Pro, proline; F, Phe, phenylalanine; W, Trp, tryptophan; Y, Tyr, tyrosine; S, Ser, serine; Q, Gln, glutamine; T, Thr, threonine; N, Asn, asparagine; D, Asp, aspartic acid; E, Glu, glutamic acid; K, Lys, lysine; R, Arg, arginine; H, His, histidine; C, Cys, cysteine.

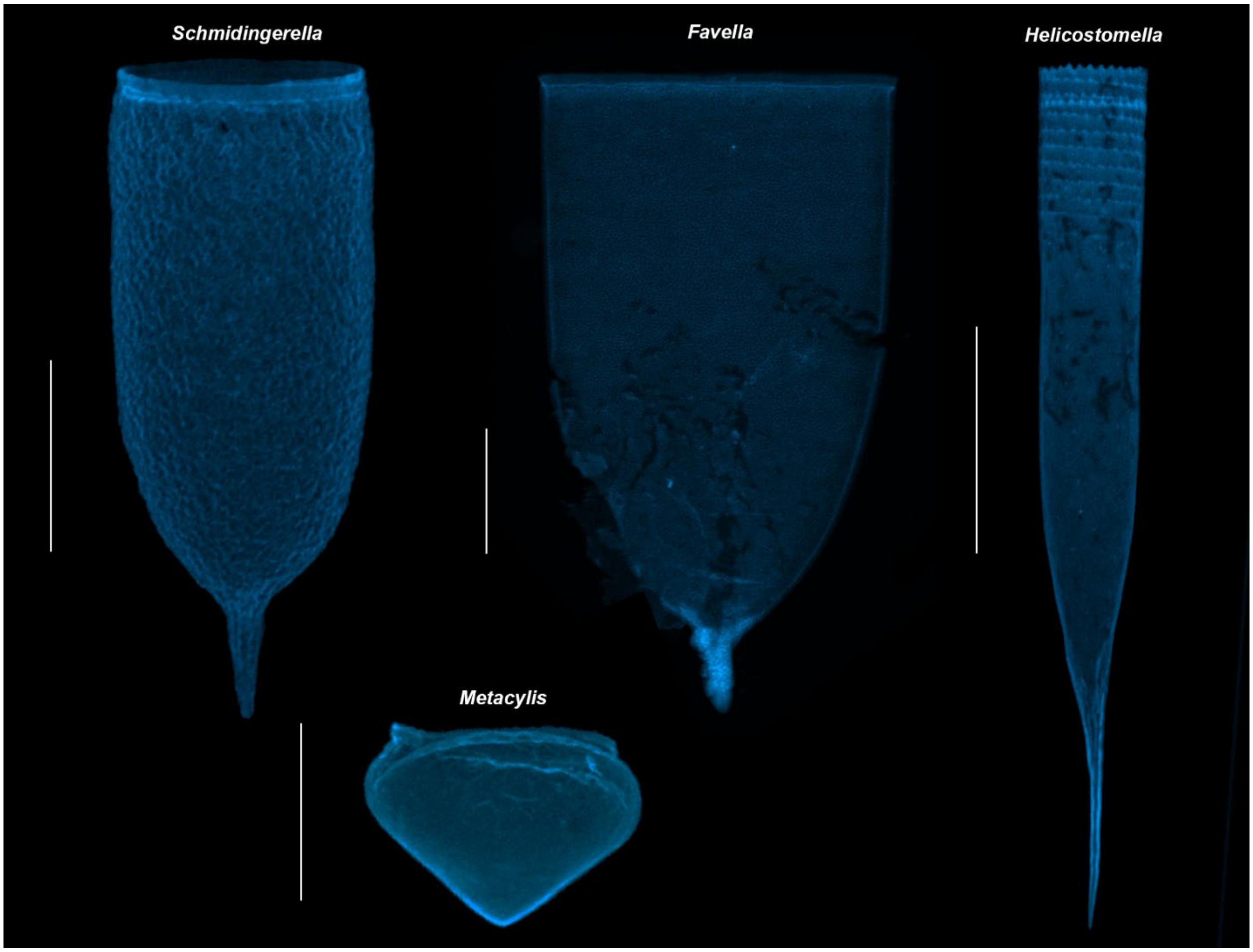


**Fig. S5. Autofluorescence of tintinnid ciliate shells excited with UV light**. The optically transparent shells of *Schmidingerella*, *Favella*, *Helicostomella*, and *Metacylis* exhibit an emission at about 465 nm after an excitation at 385 nm. The shell of *Favella* is squashed due to the pressure of the cover slip. Scale bars are 50 µm.

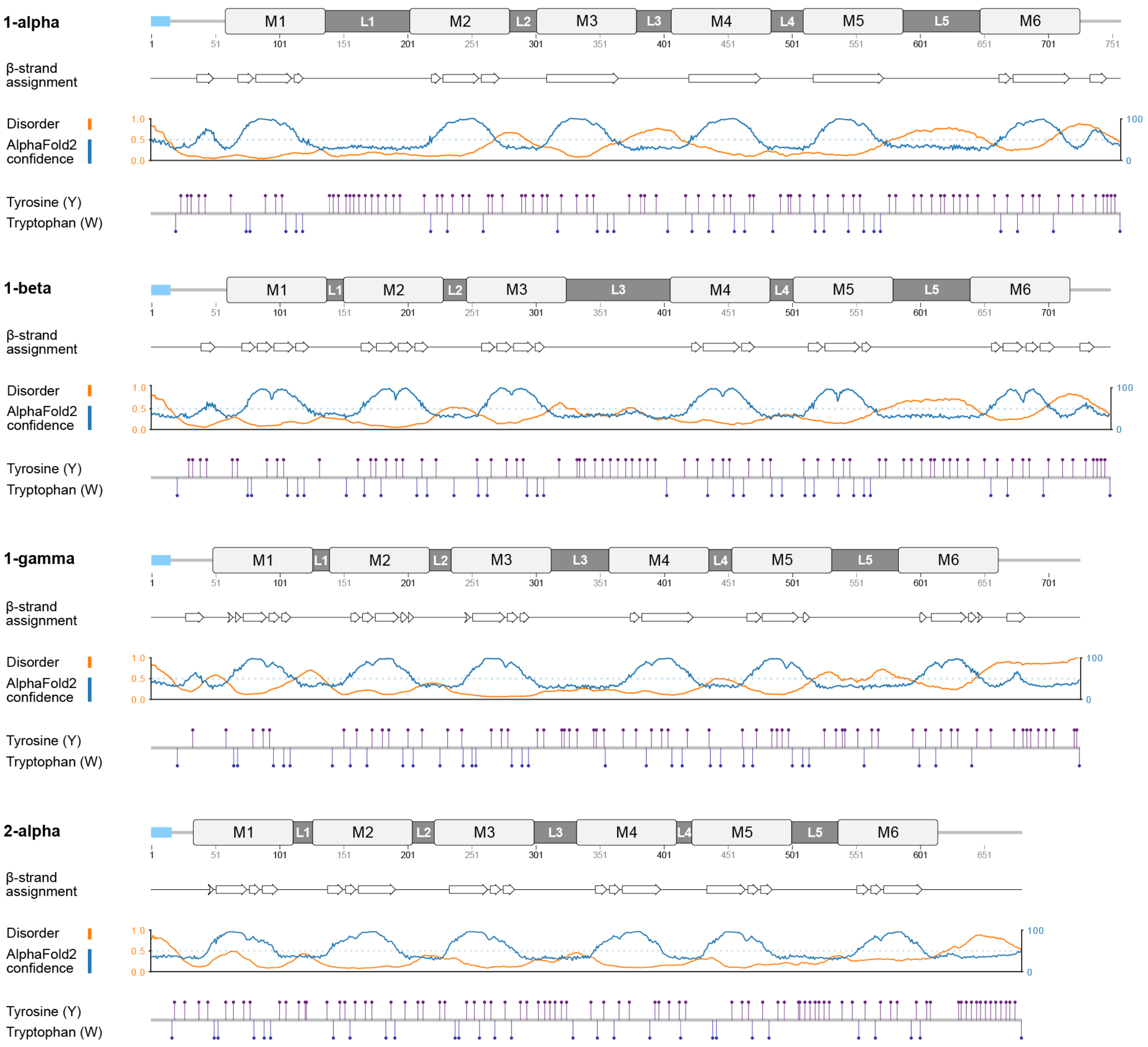


**Fig. S6. Sequence and structural characteristics for Tintinnidorin-1-alpha, beta, and gamma of *Schmidingerella* and Tintinnidorin-2-alpha of *Tintinnopsis cylindrica*.** Upper lines: Schemes of the six modules (M1–6), the connecting linkers (L1–5), and the signal peptides (light blue). Second line: Positions of beta-strands as assigned by STRIDE *(44)*. Third line: Intrinsically disordered regions (orange) as predicted by metapredict V2 (*45*) and AlphaFold2 (blue) per-residue confidence score (pLDDT, predicted local distance difference test). Disorder values (orange) higher than 0.5 (threshold = dotted line) indicate high probability for disorder. High AlphaFold2 model confidence values are generally predicted for the modules and concur with low values for disorder. Conversely, high values for disorder are mainly predicted for the linker segments, for which the model confidence values are low. Fourth line: Distributions of tyrosine (Y) and tryptophan (W) residues.

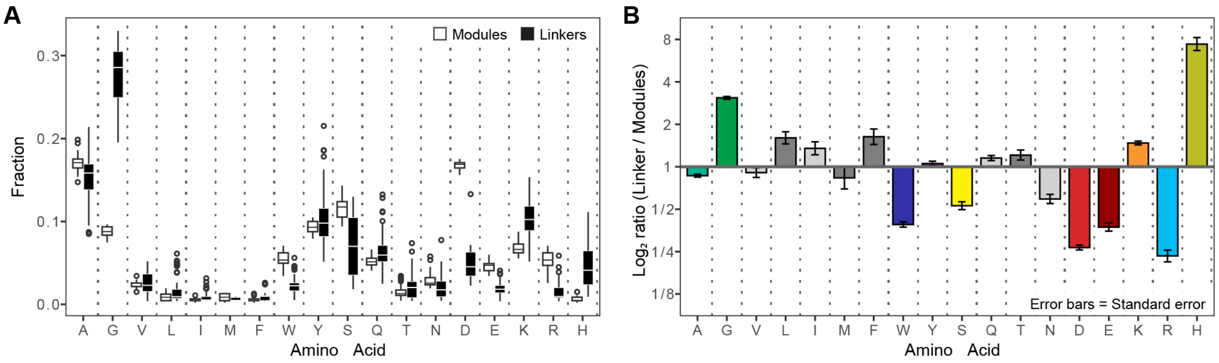


**Fig. S7. Comparisons of amino acid compositions between modules and linkers in the 78 full-length Tintinnidorin proteins. (A)** Fractions. **(B)** $\text{Log}_2$ ratios of average linker and module fractions. A, Ala, alanine; G, Gly, glycine; V, Val, valine; L, Leu, leucine; I, Ile, isoleucine; M, Met, methionine; Pro, proline; F, Phe, phenylalanine; W, Trp, tryptophan; Y, Tyr, tyrosine; S, Ser, serine; Q, Gln, glutamine; T, Thr, threonine; N, Asn, asparagine; D, Asp, aspartic acid; E, Glu, glutamic acid; K, Lys, lysine; R, Arg, arginine; H, His, histidine; Cys, cysteine.

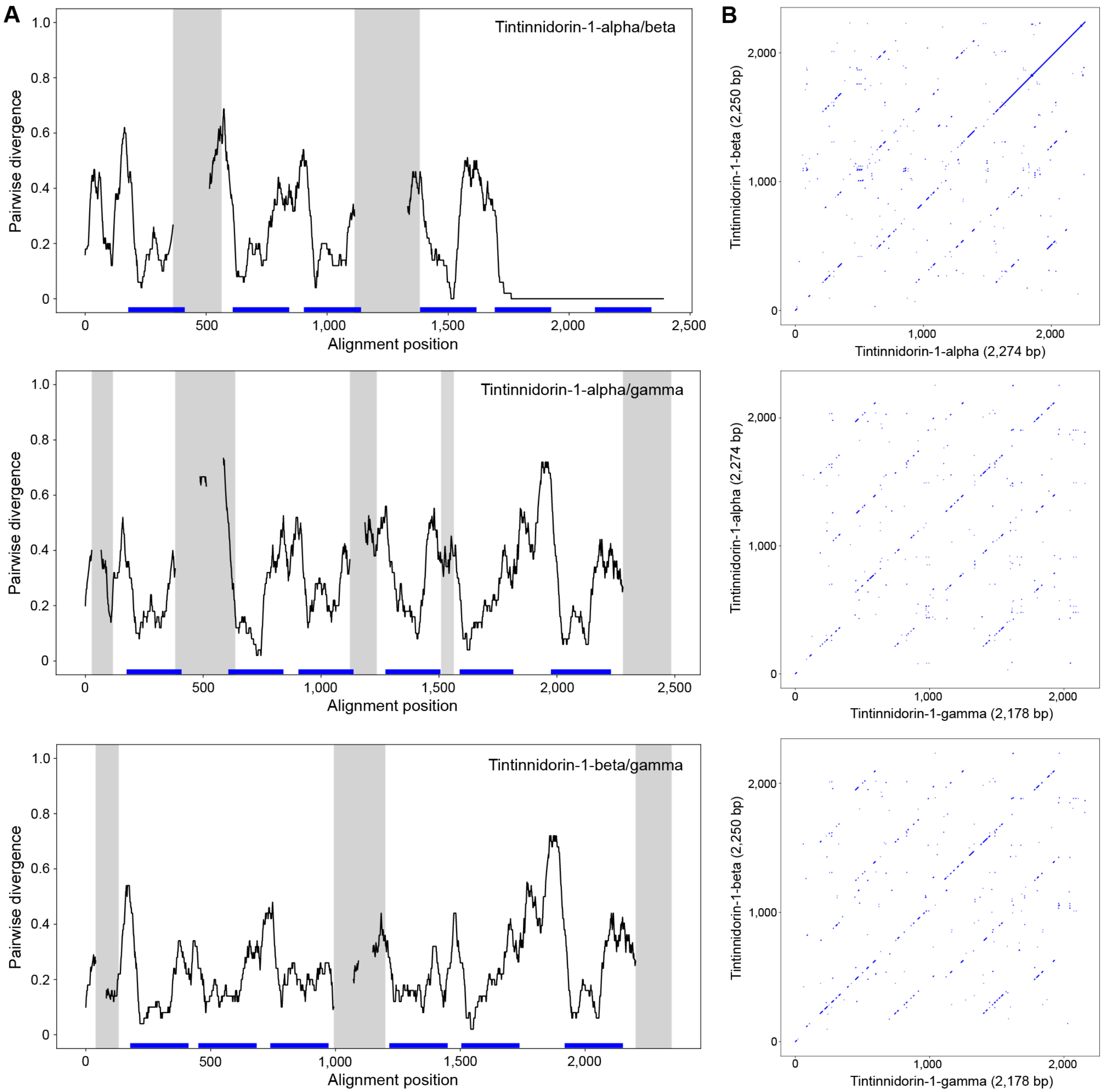


**Fig. S8. Pairwise nucleotide sequence divergence and similarity of Tintinnidorin-1-alpha, beta, and gamma. (A)** Sliding window analyses (50 base pairs with a step size of one base pair) of aligned nucleotide sequences of Tintinnidorin-1-alpha, beta, and gamma. Divergence is only partly plotted in the grey shaded alignment segments when one of the two compared sequences contains many gaps (≥ 20). Blue lines above the x-axes mark the six module positions in the alignment. **(B**) Dotplots visualizing similarity between sequence pairs. Forward matches (blue dots) were identified, using a sliding window of 10 base pairs.

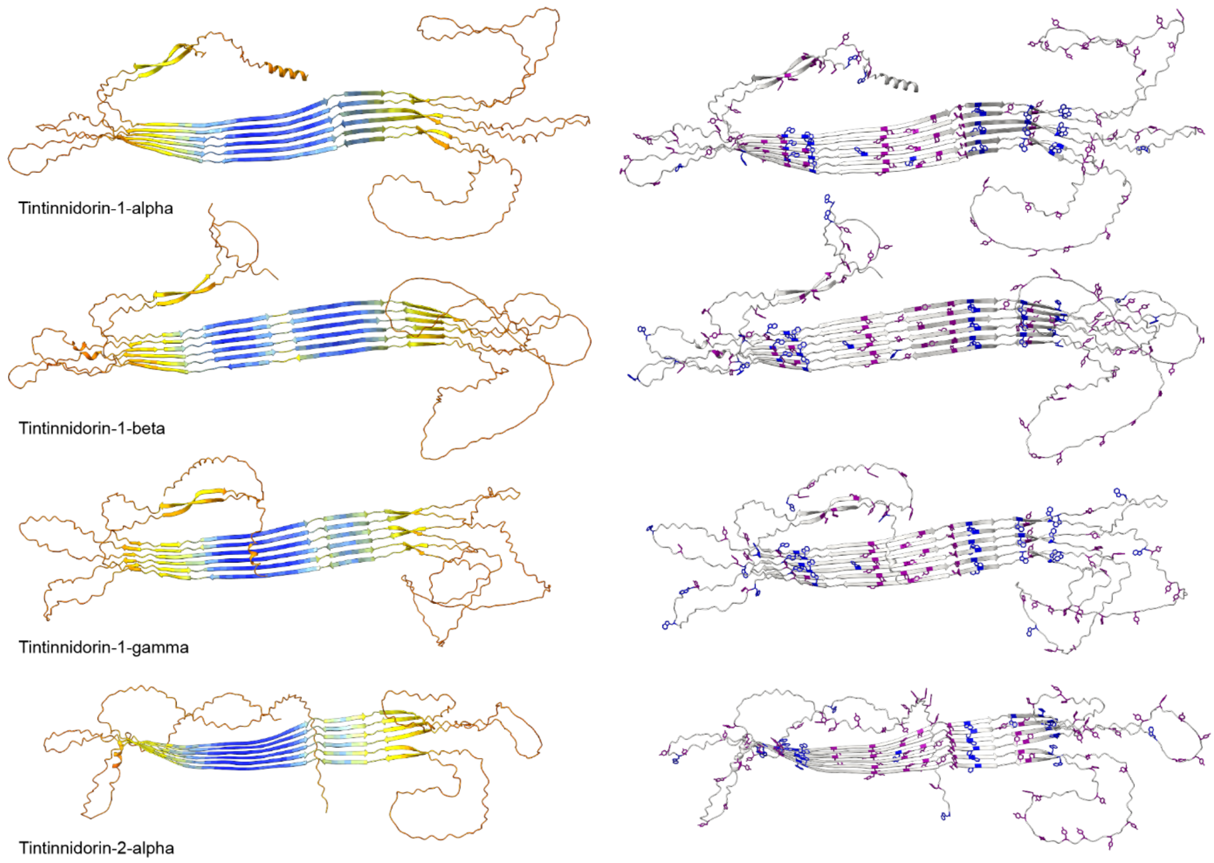


**Fig. S9. Structure models of Tintinnidorin-1-alpha, beta, and gamma from *Schmidingerella* and Tintinnidorin-2-alpha from *Tintinnopsis cylindrica* predicted by AlphaFold2.** First column: Models color-coded by per-residue scores (pLDDT, predicted local distance difference test) ranging from high (blue) to low (orange) confidence values. Second column: Distribution of tyrosine (purple) and tryptophan residues (blue).

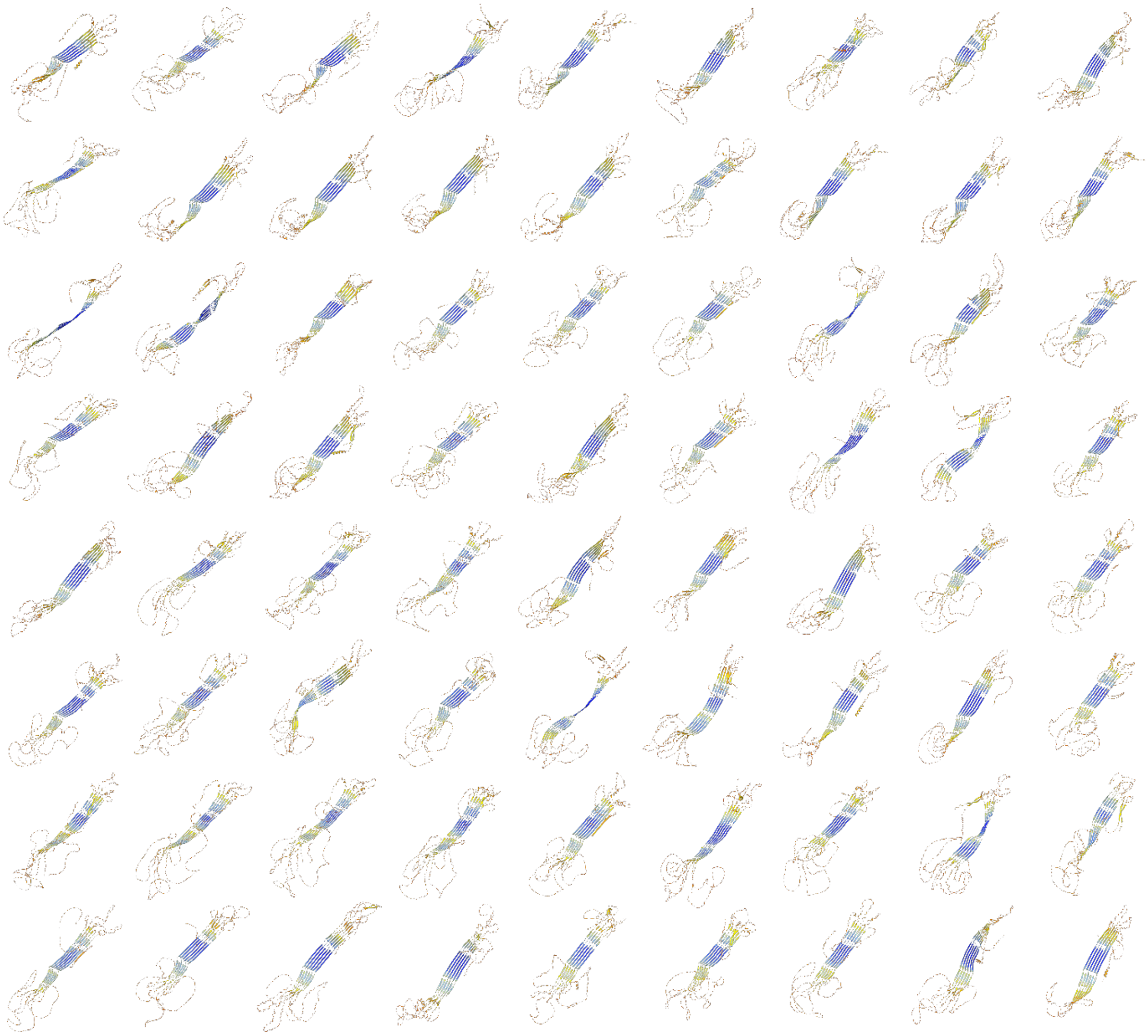

**Fig. S10. Structure models of the 72 full-length Tintinnidorin proteins extracted from the Tara Oceans database and North Pacific Eukaryotic Gene Catalog.** The models are predicted by AlphaFold2 and color-coded by per-residue scores (pLDDT, predicted local distance difference test) ranging from high (blue) to low (orange) confidence values. Consistently, the core structure segments (antiparallel beta-sheets) are folded with the highest confidence, whereas the remaining segments are mostly disordered.

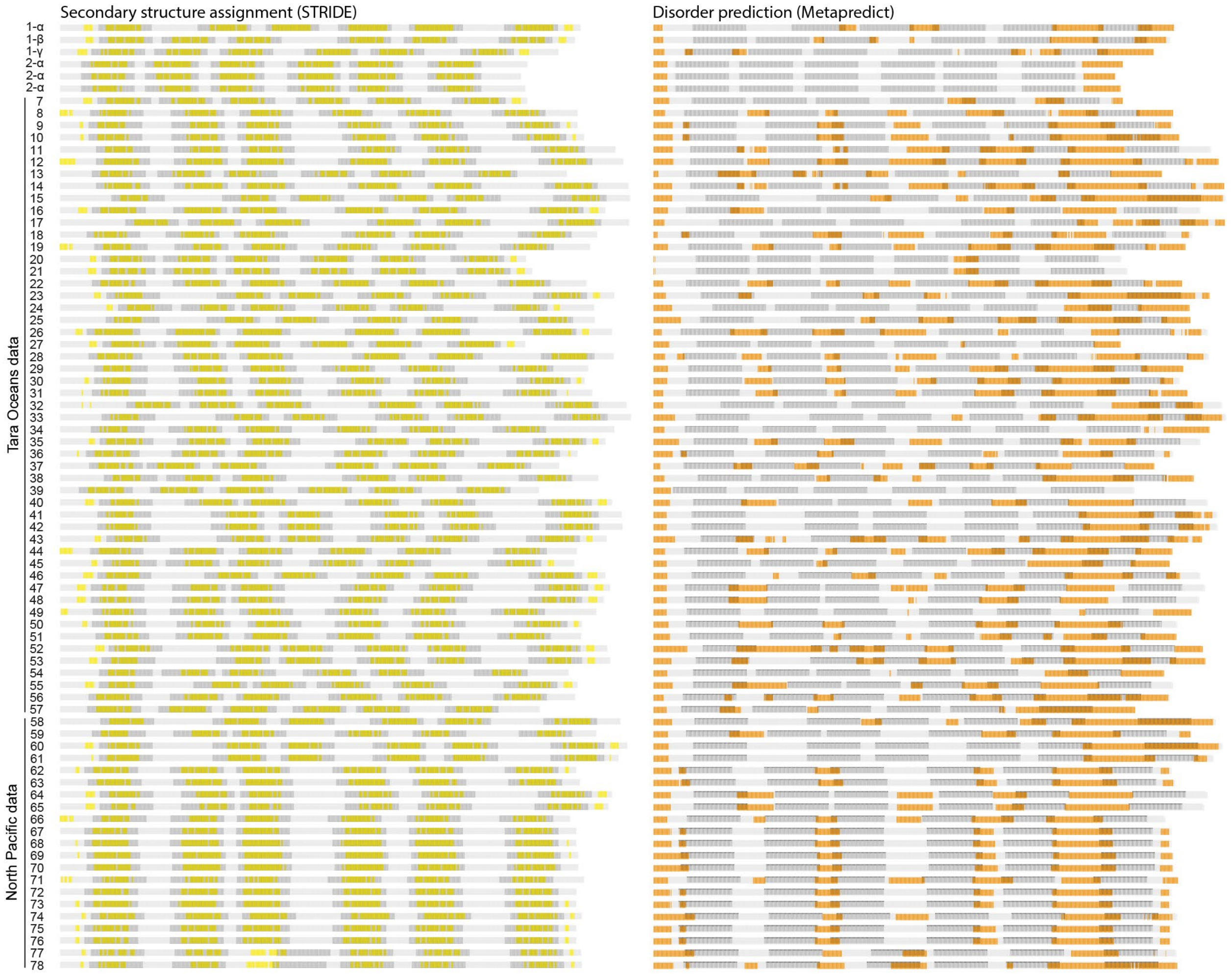


**Fig. S11. Secondary structure assignments and propensities for disorder in the 78 full-length Tintinnidorin proteins.** Congruently, beta-sheets (left column; yellow) are assigned as the main secondary structure to the six modules of each sequence by STRIDE. In contrast, propensities for disorder (right column; orange) are predicted by metapredict V2 mainly for the N- and C-terminal segments and the linkers, especially for each most C-terminal linker. The sequence numbers refer to further information in table S7. 1-α, 1-β, 1-γ, Tintinnidorin-1-alpha, beta, and gamma of *Schmidingerella*; 2-α, Tintinnidorin-2-α of *Tintinnopsis cylindrica*.
